# Integrative analysis of hexaploid wheat roots identifies signature components during iron starvation

**DOI:** 10.1101/539098

**Authors:** Gazaldeep Kaur, Vishnu Shukla, Anil Kumar, Mandeep Kaur, Parul Goel, Palvinder Singh, Anuj Shukla, Varsha Meena, Jaspreet Kaur, Jagtar Singh, Shrikant Mantri, Hatem Rouached, Ajay Kumar Pandey

## Abstract

Iron is an essential micronutrient for all organisms. In crop plants, iron deficiency can decrease crop yield significantly, however our current understanding of how major crops respond to iron deficiency remains limited. Herein, the effect of Fe deprivation at both the transcriptomic and metabolic levels in hexaploid wheat was investigated. Genome-wide gene expression reprogramming was observed in wheat roots subjected to Fe starvation, with a total of 5854 genes differential expressed. Homoeolog and subgenome specific analysis unveiled induction bias contribution from the A and B genomes. In general, the predominance of genes encoding for nicotianamine synthase, yellow stripe like transporters, metal transporters, ABC transporters and zinc-induced facilitator-like protein was noticed. Expression of genes related to the strategy-II mode of Fe uptake was predominant as well. Our transcriptomic data were in agreement with the GC-MS analysis that showed the enhanced accumulation of various metabolites such as fumarate, malonate, succinate and xylofuranose, which could be contributing to Fe-mobilization. Interestingly, Fe starvation leads to significant temporal increase of glutathione-S-transferase both at transcriptional and in enzymatic activity levels, which indicates the involvement of glutathione in response to Fe stress in wheat roots. Taken together, our result provides new insight into the wheat response to Fe starvation at molecular level and lays foundation to design new strategies for the improvement of Fe nutrition in crops.

## Introduction

Iron (Fe) is among the essential micronutrients in plants that participate as catalytic cofactors in several key processes including photosynthesis, respiration, chlorophyll biosynthesis, and nitrogen fixation (Kim and Rees, 1992; Morrissey and Guerinot, 2009; Li et al., 2017). The bioavailability of Fe in the soil is strongly dependant on its solubility, aerobic and calcareous soil condition, pH, and the presence of natural ligands secreted by plant roots (Marschner H., 1995; Morrissey and Guerinot, 2009; Thomine et al., 2013). To circumvent the above challenges for Fe uptake by the roots, plants have adapted two modes of uptake strategies. Strategy-I, mostly predominant in Eudicot species, primarily relies on the enrichment of rhizospheric regions with protons (H^+^) and other reducing agents (Hell and Stephan, 2003; Santi et al., 2005; Santi and Schmidt 2009; Kobayashi and Nishizawa 2012). In contrast, graminaceous species follow the Strategy-II mode of uptake, which involves the transport of Fe in the complex-chelated form (Mori et al., 1999; Kobayashi and Nishizawa, 2012; Connorton et al., 2017). The primary components involved in chelation are the phytosiderophores (PS) secreted by plant cells in the rhizospheric region mainly by efflux transporters (Morrissey and Gueirnot, 2009; Kobayashi and Nishizawa, 2012). One of the main components of these secreted siderophores involved in Fe chelation is referred to as mugineic acids (MAs). The transporters involved in the secretion of the MAs are identified as transporter <u>o</u>f <u>m</u>ugineic (TOM) acid proteins (Nozoye et al., 2011). The complex chelated form of Fe^3+^-PS is taken up by the specific root transporter referred to as yellow stripe-like transporter proteins (YSL) (Curie et al., 2001; Gross et al., 2003, Yordem et al., 2011). Subsequently, Fe is transported in the plant organelles by multiple partners including specialized long distance, tissue-specific transporters as reported in monocots (e.g rice, maize) and eudicots (e.g. Arabidopsis) (Kim et al., 2006; Waters et al., 2011). In other monocots such as maize and rice, the presence of genetic components for both Strategy-I and II were reported (Ishimaru et al., 2006; Inoue et al., 2009; Lee et al., 2009; Zanin et al., 2017). At the metabolome level, analytical approaches utilizing GC-MC and LC-MS were utilized to study the components of Fe starvation in plants (Palmer et al., 2014; Kabir et al., 2013). However, the molecular and metabolic activity predominant for Fe uptake by the roots of hexaploid wheat still remains to be elucidated.

Microarrays have been successfully used to investigate the global transcriptional changes in Arabidopsis plants grown in Fe starved conditions (Thimm et al., 2001; Buckhout et al., 2009; Yang et al., 2010). Significant changes in the expression pattern of the genes primarily involved in Strategy-II mode of uptake were observed in roots of maize and rice subjected to Fe-deficiency stress (Li et al., 2014; Quinet et al., 2012; Kobayashi et al., 2014; Bashir et al., 2014). The transcriptome analysis of Fe-starved plants is characterised by an important change in gene expression of several transcriptional regulators such as transcription factors (TFs) (Colangelo et al., 2004; Kobayashi et al., 2009; Long et al., 2010; Li et al., 2016; Connorton et al., 2017) or key genes related to phytohormone homeostasis (Schmidt et al., 2000; Hindt and Guerinot, 2012). Fe starvation also leads to changes in the abundance of transcripts related to plant metabolism and genes involved in signalling pathways that modulate nutrient uptake. Additionally, genes involved in ethylene/auxin signalling or linked with certain other macronutrients like nitrogen, sulphur and phosphorus are also significantly expressed (Romera et al., 1994; Zheng et al., 2009; Romera et al., 2011; Borlotti et al., 2012; Zuchi et al., 2015; Lin et al., 2016; Zanin et al., 2017; Garnica et al., 2018). Therefore, transcriptome analysis is a powerful approach to help understand the network involved in plants response to Fe.

Hexaploid wheat is an important crop that is also a good, affordable source of nutrition. Fe deficiency strongly affects the growth of cereal crops and productivity (Yousfi et al., 2009; Bocchini et al., 2015). The development of transcriptome technology such as RNA sequencing-RNAseq and the availability of genome sequence for hexaploid wheat (genome A, B, D) (International Wheat Genome Consortium, 2014) combined with metabolomic approaches will help improve our knowledge of wheat response to nutritional stress (e.g. –Fe) (Borrill et al., 2018). Our current knowledge on how the wheat genome responds to Fe limiting condition remains obscure and unanswered. In addition, how the different genomes could impact the homoeolog based expression of the transcripts, categorically under different stress conditions, remains to be explored (Ramirez-Gonzalez et al., 2018). In the current study, RNA-seq based approach was utilized to address the molecular components involved in wheat roots during Fe starved condition. Further, genome-based expression study was performed that provided preliminary clue on coordinated expression of the homoeologs. Metabolic profiling result support the data from the transcriptome study and pinpoint the role of glutathione mediated response. Collectively our results provide the first insight on molecular and biochemical responses of hexaploid wheat under Fe starvation.

## Materials and methods

### Plant materials, starvation conditions and plant sampling

Hexaploid bread wheat variety ‘C-306’ was used for the experimental purpose. Wheat grains were subjected to stratification in dark at 4 °C for overnight and were allowed to germinate for 5 days on the Petri plates containing 2-3 layers of wet Whatman filter paper. The endosperms were removed from the developing seedlings once they started browning. Subsequently, the seedlings were transferred in PhytaBox™ and grown for 7 days in Hoagland’s nutrient solution. A total of forty-eight seedlings were kept equally in four PhytaBoxes, used for each treatment. After 7 days, nutrient solutions were replaced on the basis of different treatments. For Fe starvation (–Fe) 2 µM Fe (III) EDTA was used as the Fe source. For control plants (+Fe) concentrations of nutrients were unchanged in above-mentioned Hoagland’s solution containing (20 µM Fe (III) EDTA). Treated seedlings were grown in the described medium for 20 days in a growth chamber set at 20±1 °C, 50-70% relative humidity and photon rate of 300 μmol quanta m^−2^ s^−1^ with 16 h day/8 h night cycle. For sampling, roots and shoots were collected at different time points after starvation (5, 10, 15 and 20 days). On the basis of distinct phenotype, samples collected at 20 days after starvation (DAS) were used for transcriptome analysis. A total of four biological replicates (each containing 10-12 seedlings) were used. Subsequently, RNA extractions were performed independently for each of the pools. Prior to RNA sequencing, quality of RNA was checked and extractions derived from two replicates were pooled together thereby generating two experimental samples for each of the respective conditions. Remaining samples were snap frozen in liquid nitrogen and stored at −80°C. To distinctively observe primary root and 1^st^ order lateral root, individual plants were moved onto a 150 mm wide petri plate filled with distilled water and characteristics were manually examined. To ascertain root characteristics, 6-8 wheat seedlings were used for each of the treatments with two experimental replicates.

### RNA extraction and Illumina sequencing

Total RNA was extracted from the treated root samples along with the control by Trizol (Invitrogen, ThermoFisher, USA) as per the manufacturer’s instruction. The quality and quantity were checked on the 1% denaturing RNA agarose gel and Nanodrop respectively. Subsequently, all the RNA used for library preparations were checked for their RNA integrity number ≥ 8.5 using Bioanalyzer (Agilent, USA). The quality control (QC) passed RNA samples were then processed for library preparation. The Paired-ended (PE) libraries were prepared from the total RNA using Illumina TruSeq stranded mRNA library prep kit as per the instructions (Illumina Inc., USA). The generated mean of the libraries fragment size distribution was 559 bp, 584 bp, 546 bp and 604 bp for the samples. The generated libraries were sequenced on the NextSeq 500 using 2 × 150 bp chemistry. The raw reads were processed further before the sorted high-quality reads were mapped to the reference genome.

### RNA-Seq analysis

Adapter clipping and quality trimming of the raw reads were performed using Trimmomatic-0.35. The sequenced raw reads were processed to obtain high quality clean reads. Ambiguous reads (reads with unknown nucleotides “N” larger than 5%) and low-quality sequences (more than 10% quality threshold QV<20 phred score) were removed. A minimum length of 100 nts after trimming was applied. Finally, high-quality (QV>20), paired-end reads were used for reference based read mapping. The genome of *Triticum aestivum* L. was taken as reference for analysis. The Gene Feature Format files were downloaded from Ensembl Plants (TGACv1.37). The reads were mapped to the reference genome using TopHat v2.1.1 with default parameters. Cufflinks v2.2.1 program assembled the transcriptome data from RNA-seq data and quantified their expression. Mapped reads were subjected to Cufflinks, followed by Cuffmerge and Cuffdiff. Log2 Fold Change (FC) values greater than one was considered up-regulated whereas less than 1 were considered as down-regulated. These genes were further categorized on the basis of statistical significance (p<0.05) and the False Discovery Rate (FDR 0.05) after Benjamin-Hochberg corrections for multiple testing for their significant expression.

Comprehensive gene annotation of wheat sequences was done using KOBAS 3.0 (Xie et al., 2011) annotate module by alignment with Rice sequences (BLASTP, cutoff 1e-5). MapMan was used to visualize the pathways involving wheat differentially expressed genes (DEGs). Pathway enrichment analysis was performed using KOBAS standalone tool. MeV was used to construct heatmaps for selected DEGs using the normalized expression values of genes. The data generated from this study has been deposited in the NCBI Sequence Read Archive (SRA) database and is accessible with the submission ID-SUB5206887 and BioProjectID-PRJNA529036.

### Gene annotation filtering and functional enrichment analysis

Significant sets of DEGs under iron starvation were further mapped using GO and MapMan. The GO annotation was downloaded from ensembl plants (https://plants.ensembl.org/biomart/martview). Mercator (Lohse et al., 2014) was used to build MapMan mapping file for TGACv1 sequences and DEGs were visualized in MapMan v3.1.1 tool (Thimm et al., 2004). For functional categorization of DEGs that were positively and negatively correlated with iron starvation, BINGO version 3.0.3 (Maere et al., 2005) was used to perform GO enrichment analysis with hypergeometric test and significant terms with an FDR value below 0.05 were considered. For gene ontology mapping, GO_full.obo ontology file was downloaded from GO consortium. MapMan classification was used to categorize DEGs into transcriptional factors. The results were visualized as network using EnrichmentMap version 2.2.1 (Merico et al., 2010) and gene expression overview in various pathways were visualized in MapMan tool.

### Homoeolog specific expression analysis for genome bias

To identify wheat homoeologous triplets; **e**nsembl biomart TGACv1 was used to extract all possible homoeologous relations. This led to 86830 pairwise homoeologous relations. An in-house script was used to select only the triplets where contribution from each genome was 1:1:1 (A:B:D), thus generating 16850 triplets. Further, triplets resulting from potential translocation events were not considered, i.e., only homoeolog triplets from same chromosome (eg., 1A, 1B, 1D triplet is accepted, whereas, 2A, 3B, 3D is rejected) were taken for analysis. Finally, 15604 triplets (15604*3 = 46812 genes) were used for studying genome induction biasness in response to iron stress. Paired end reads were aligned to the reference (selected scaffolds from genome that harbour 15604 triplets) using TopHat v2.1.1 with a specific argument (--b2-very-sensitive) (Powell et al., 2017), which leads to more stringent alignments as required for homoeologs. The Cufflinks pipeline was used to obtain FPKM values and differentially expressed genes.

Further, relative abundance levels and expression bias for homoeologs was studied by considering the homoeologous gene triplets with expression of FPKM ≥ 1 each in both the control as well as Fe starved conditions. For this, the normalised relative expression for each homoeolog within a triad was calculated. For example, the relative expression from A will be represented as:

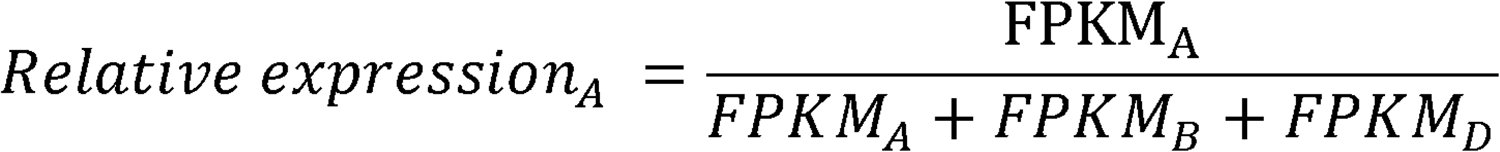

Thus, relative expression levels of A, B and D homoeologs within each triad were calculated similarly. Seven homoeolog expression bias categories were defined as described earlier by Ramirez-Gonzalez et al., (2018). In total, seven categories defined as one balanced category and six unbalanced homoeolog-suppressed/homoeolog-dominant (from either of the genomes) were listed for the ideal relative expression of A, B and D. Eucledian distance (using cdist function from rdist package, R3.3.2) was calculated between normalised relative expression for each triad and the seven ideal categories. The shortest distance was used as a deciding factor to group the triads into the seven respective categories.

### Quantitative real time-PCR (qRT-PCR)

For validation of the gene expression, qRT-PCR analysis was performed. Total RNA (2 µg) isolated from the above experiments were used to validate the expression by using qRT-PCR method. Genomic DNA present in trace amount was removed by DNaseI treatment using a Turbo DNA-free kit (Invitrogen, ThermoFisher, USA). Further, cDNA was synthesized from two micrograms of DNA-free RNA using Superscript III first strand (Invitrogen, ThermoFisher, USA) with random hexamer primers following the manufacturer’s guidelines. For qPCR reaction, gene-specific primers of each gene were designed from conserved regions of all three homeolog sequences (Supplementary Table S1). qRT-PCR was performed using QuantiTect SYBR Green RT-PCR mastermix (Qiagen, USA) with programs recommended by the manufacturer in the ABI 7700 sequence detector (Applied Biosystems, USA). ADP-ribosylation factor (ARF) and Actin were used as internal controls. Two independent experimental replicates with four technical replicates were performed for each sample. The relative amount of gene expression was calculated using 2^− ΔΔCT^ method.

### Metabolite extraction and GC-MS profiling

Extraction of total metabolites was performed similarly as previously described (Wang et al., 2018). Roots of plants grown under –Fe and +Fe were sampled in triplicate manner and dried for 1 week. Each of 50 mg crushed samples was extracted in a mixture of solvents (precooled; 300 µl methanol, 30 µl 2 mg ml^−1^ nonadecanoic acid methylester and 30 µl 0.2 mg ml^−1^ sorbitol) for 15 minutes (70°C, 1000rpm). Further, at room temperature 200 µl chloroform was added and shaken for 5 min (37°C, 1000rpm). To obtain phase separation, 400 µl H_2_O was added to each sample, vortexed and centrifuged (10 min, 13,000 rpm). From the upper polar phase approximately 200 µl was finally aliquoted for complete drying.

For GC-MS analysis, metabolites were subjected to methoxyamination and trimethylsilylation. Dried polar phase were shaken for 1.5 hr at 30°C in 40 µl of MeOX (40 mg ml^−1^ methoxyaminhydrochloride in pyridine) followed by 30 min shaking at 37°C in 80 µl BSTFA mixture (70 µl N,O-bis(trimethylsilyl)trifluoroacetamide + 10 µl alkane mix). The derivatized metabolites were subjected to GC-MS analysis (Agilent technologies 7890, USA) coupled with mass spectrometry. Measurement from an injection volume of 1 µl was taken in split-less mode in DB-5 column (30 m × 0.25 mm, 0.25 µm film thickness, Agilent) using helium as carrier gas. Metabolites were separated as described by Wagner et al. (2013). Qualitative analysis of chromatograms was performed in MassHunter Qualitative analysis Sp1 workstation (Agilent, USA). Identification and annotation of each compound was supervised manually using AMDIS software and NIST08 database (http://www.nist.gov/srd/mslist.html). Data were normalized to sample weight and internal control (sorbitol). Statistical analysis was performed as described earlier (Quanbeck et al., 2012). Log2 ratio of metabolite abundances in –Fe was plotted against +Fe. Delta method approximation was used to calculate standard errors (se) of log-ratio, se log-ratio = 1/ln 2√[(SE_T_ /T)^2^ + (SE_C_ /C)^2^], where SE_T_ and SE_C_ are standard errors of average –Fe and +Fe metabolite abundances.

### Measurement of glutathione-S-transferase (GST) activity and iron mobilization assay for PS estimation

Activity measurement of GST was performed in the wheat roots subjected to –Fe for 10, 15 and 20 days of treatment along with the control plants (no stress) as mentioned in plant materials. The estimation was done using Glutathione-S-transferases assay kit (Sigma, USA). Briefly, 100 mg of tissue was used for total protein extraction. Equal amount of total protein (25 μg) was used as a source of enzyme and 1-Chloro-2,4-dinitrobenzene (DNB) was used as a substrate. The resulting GS-DNB conjugate was measured at 340 nm wavelength during the time course of the reaction. The direct increase in absorption was measured and GST activity was calculated as described in the manufacturer instruction kit.

For PS release, iron mobilization assay was performed as described earlier (Takagi et al., 1976). Briefly, 10 seedlings (each for –Fe and +Fe for 10 days treatment) were used for the experiment. After treatment the seedlings were subjected to PS release after 2 hours of onset of light period in aerobic condition for 3 hrs in 20 ml deionised water with 10 mgl^−1^ Micropur (Katadyn, Switzerland). Next, 2 ml of freshly precipitated solution of Fe(OH)_3_ and 0.5 ml of 0.5 M Na acetate buffer (pH 5.6) was added to 8 ml of the collection solution. The solution was shaken for 2 hrs and subsequently was filtered (Whatman #1) into 0.2 ml of 6NHCl. Ferric iron was reduced by addition of 0.5 ml of 8% hydroxylamine-hydrochloride and heating to 60 °C for 20 min. The total concentration of ferrous iron was calculated by measuring absorbance at 562 nm after adding 0.2 ml of 0.25%ferrozine and 1 ml of 2 M Na-acetate buffer (pH 4.7).

### Elemental analysis using inductive coupled plasma-mass spectroscopy and nitrate estimation

Elemental analysis in roots and shoots was performed using Inductive Coupled Plasma-MS (ICP-MS). Metal analysis was performed as described earlier (Bhati et al., 2016; Aggarwal, 2018). Briefly, the samples were ground to fine powder and subsequently subjected to the microwave-digested with HNO_3_ as described earlier (SuraPure™, Merck). Respective metal standards were also prepared for analysis. Three independent replicates from the experiments were used for metal analysis.

Nitrate content in wheat roots was measured according to method described previously (Cataldo et al., 1975). Briefly, 1 g of fresh tissue was homogenized in 6 ml of deionised water and centrifuged at 30,000g for 15 min. The 100 µl of supernatant was added to 400 µl of salicylic acid (w/v dissolved in conc. H_2_SO_4_). After mixing the reaction was kept at room temperature for 20 min. 2N NaOH (9.5ml) was then added slowly to raise the pH above 12. The samples were allowed to cool and readings were taken at 410nm in spectrophotometer.

## Results

### Fe starvation affects wheat growth capacity and nutrient uptake

Fe starvation is known to affect plant growth capacity. In order to determine the effect of Fe starvation, one-week old wheat seedlings, grown on complete medium (presence of Fe) were transferred to Fe starvation media for additional days. After 20 days of starvation (DAS) the wheat seedlings started showing strong phenotypic symptoms including visible chlorosis and therefore detailed study was performed for this specific time point. In response to Fe starvation, plants showed decrease in the shoot biomass with an enhanced chlorosis phenotype and shortening of the root system compared to control wheat seedlings (Fig. 1A and B). Roots of Fe-starved wheat seedlings showed decrease in number of lateral roots and significant reduction in the primary root length in comparison to control plants (Fig. 1C and D). Earlier studies have suggested that changes in the root system and Fe supply not only affect the Fe accumulation capacity but also impacts the uptake of other nutrients such as Zn, Cd etc. (Sperotto et al., 2012; Shukla et al., 2017). Therefore, effect of Fe starvation stress on the uptake of Zn, Mn, Cu, and Mg in wheat undergoing Fe stress was studied (Table 1). Out data indicated increased uptake of nutrient elements such as Zn, Mn, Cu, and Mg in roots but accumulation in shoots was either unaltered or decreased. This further support the importance to study root response to Fe starvation to gain insight on the molecular mechanism evolved by wheat to cope with nutritional stress.

**Table 1:** Metal concentration (mg/g of tissue dry weight) in roots and shoots of wheat seedlings subjected to –Fe stress.

| *Treatments* | ROOTS |  |  |  |  | SHOOTS |  |  |  |  |
| --- | --- | --- | --- | --- | --- | --- | --- | --- | --- | --- |
|  | Fe | Zn | Mn | Mg | Cu | Fe | Zn | Mn | Mg | Cu |
| **+Fe+P** | 106.73<br>± 14.2 | 14.6 ±<br>3.2 | 15.8 ±<br>2.0 | 542 ±<br>70 | 7.4 ±<br>1.1 | 72.9 ±<br>1.7 | 26.8 ±<br>.09 | 17.4 ±<br>.05 | 1031 ±<br>2.3 | 10.4 ±<br>0.6 |
| **–Fe +P** | 66.8 ±<br>17 | 40.4 ±<br>12 | 38.2 ±<br>8.9 | 912 ±<br>213 | 17.9 ±<br>3.7 | 28.6 ±<br>1.6 | 22.3 ±<br>1.05 | 22.1 ±<br>1.9 | 1013 ±<br>48 | 5.1 ±<br>0.3 |

**Fig. 1.**
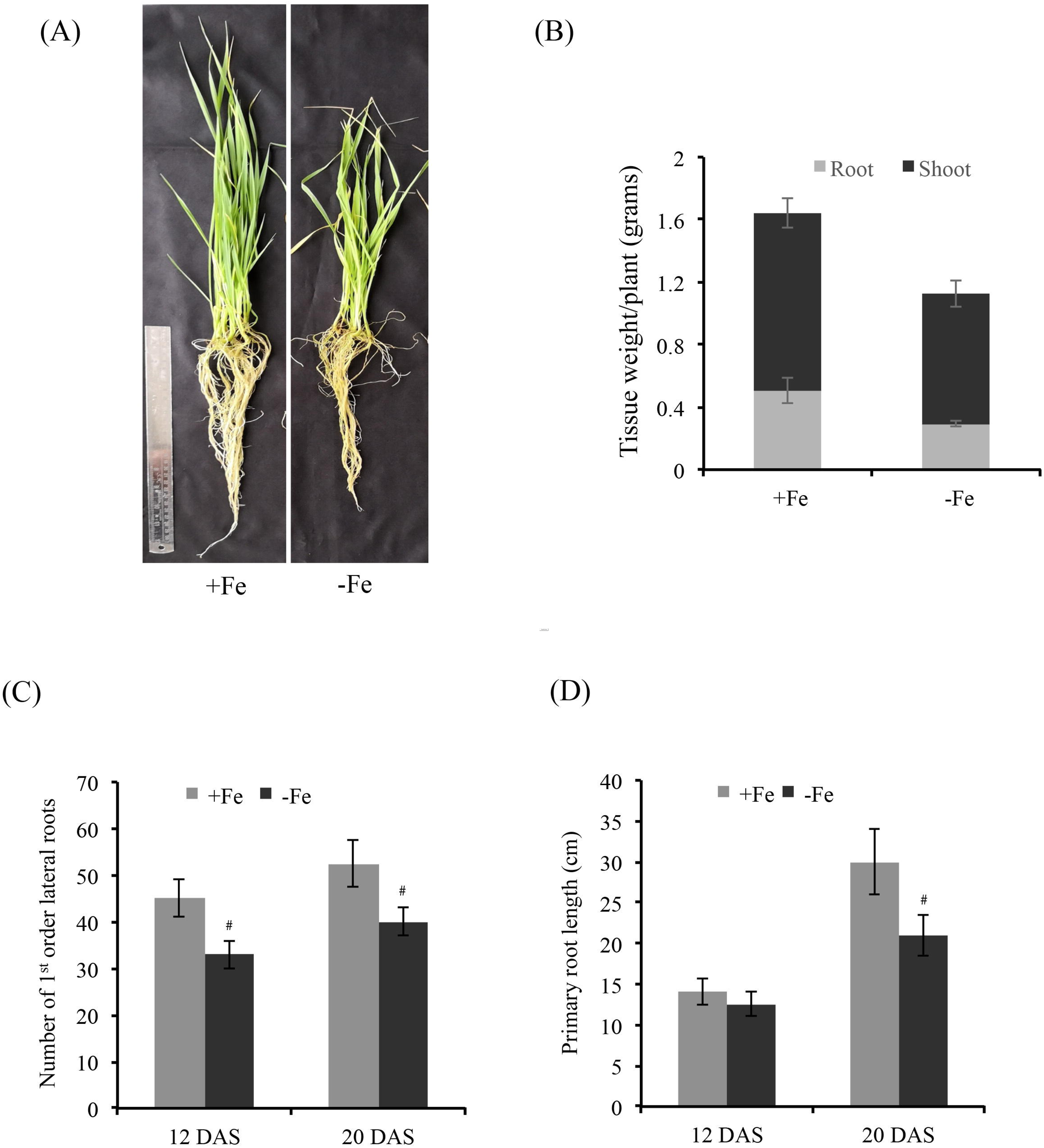
Effect of Fe-starvation (–Fe) on the growth parameters of wheat seedlings post 20 days after starvation. (A) Phenotype of wheat seedlings exposed to Fe starvation. (B) Total biomass of roots and shoots of wheat seedlings after 20 DAS. 12-15 seedlings were collected for calculating the fresh tissue weight (in grams). (C) Number of first order lateral roots in roots subjected to –Fe condition and control plants (+Fe). (D) Primary root length of wheat roots. 10-12 seedlings were used for measuring the total primary root length of wheat seedlings under –Fe and +Fe condition. # indicate significant difference at p<0.05.

### Differential expression analysis, homoeolog (A, B and D) induction and expression bias during –Fe response

The effect of Fe starvation on the wheat root transcriptome has not been investigated till date. To perform this study, RNAseq technology was used to identify the changes in the transcripts of wheat roots, where plants were grown in presence or absence of optimal Fe (20 DAS). Our analysis resulted in 87 million quality filtered reads, with an average of nearly 22 million reads from each sample (more than 87% reads had a quality score greater than Q30). Filtered reads from the four libraries had a mapping rate ranging from 81.7% to 85.4% when mapped against release-37 of the wheat genome using TopHat (Supplementary Table S2). As quality check, a strong correlation within the two biological replicates from each condition was observed, while a clear variation was seen between the two conditions (Fig. 2A). We thereafter analysed the expression values as FPKMs (Fragment Per Kilobase of transcript per Million mapped reads), calculated by using Cufflinks software. Differentially expressed genes (DEGs) were then identified by calculating logFC (log2 fold change) and performing statistical tests between FPKM values from control and stressed samples using CuffDiff. 50610 genes with an FPKM of greater than or equal to 1 in at least one of the two conditions were considered to be “expressed transcripts”. In all, 7221 genes had logFC > 0 and 8010 had logFC< 0 (Fig. 2B). On setting up a criterion of logFC of more than 1 for up-regulated genes, and that of less than −1 for down-regulated genes and an FDR < 0.05, a total of 3478 genes were highly expressed, whereas, 2376 were down-regulated under –Fe condition in wheat roots (Fig. 2C). Interestingly, 45 genes were also induced exclusively during starvation condition when compared to the control samples (Fig. 2C, Supplementary Table S3).

**Fig. 2.**
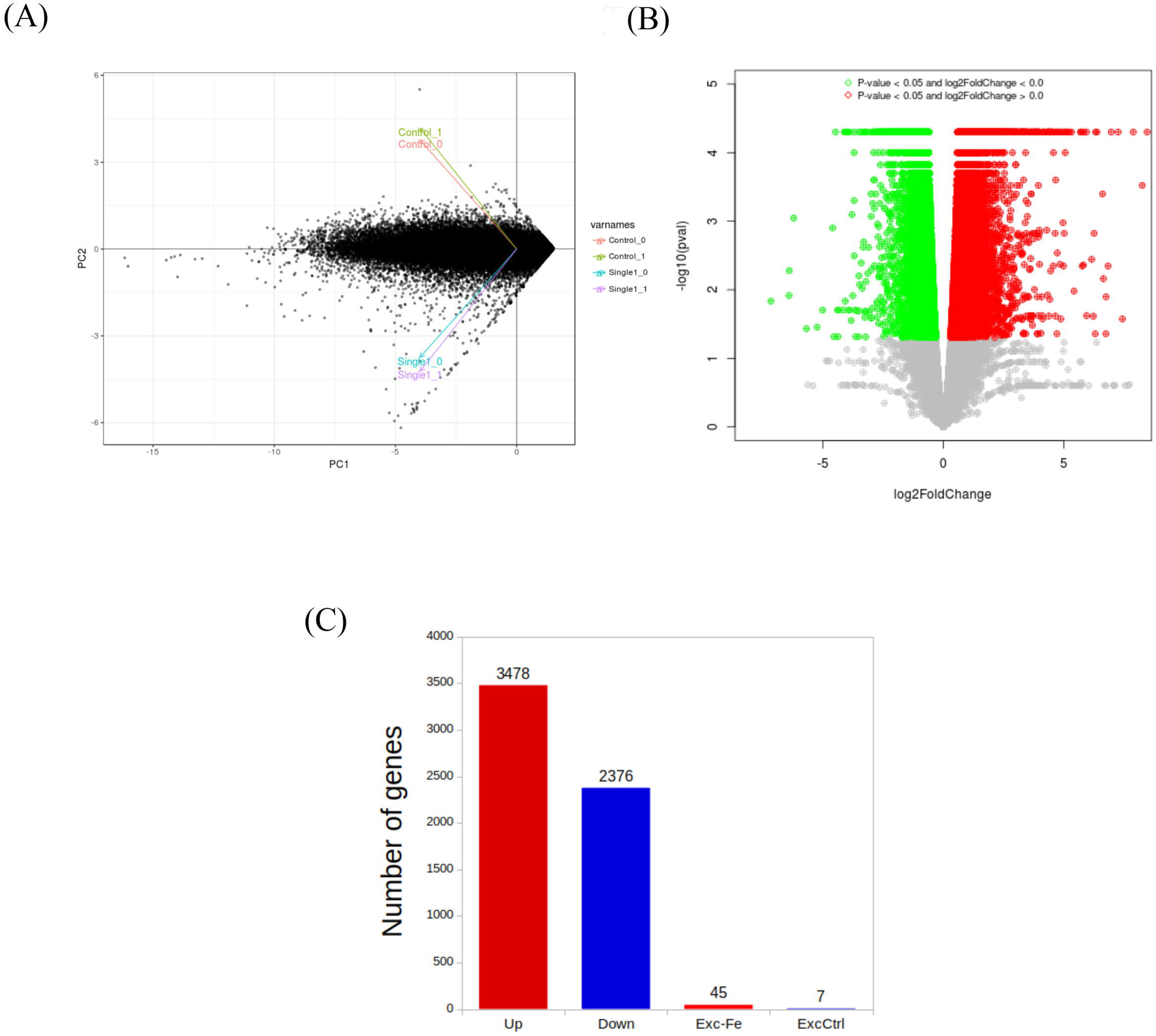
Analysis of RNAseq data from the wheat roots during Fe starvation. (A) Principal component analysis of samples from control (Control_0, Control_1) and –Fe (Single1_0, Single1_1) conditions. (B) Volcano plot of DEGs, the x-axis shows the log2 fold change difference in the expression of genes in iron starved condition w.r.t. control, and the y-axis indicates the negative log of p-value (pval) for the differences in expression. Genes without significant differences are indicated by grey dots. Significant genes with logFC > 0 are represented by red dots, and those with logFC < 0 are represented by green dots in the scatter plot. (C) Number of DEGs in –Fe. Up: Upregulated under –Fe (logFC > 1), Down: Downregulated under –Fe (logFC < −1), Exc–Fe: exclusively expressed in –Fe, ExcCtrl: Exclusively expressed in control condition.

Our data allowed us to analyse the chromosomal distribution of the DEGs under the – Fe condition. While all chromosomes contributed the DEGs, the highest number of genes was mapped on chromosome 2 of the wheat genome (Fig. 3A). Equal representation of transcripts was observed for the chromosome 7 and 5. The remaining chromosomes, 1, 3, 4 and 6 showed 15, 13, 12 and 10% distribution of DEGs, respectively. In polyploidy crops like wheat, homoeolog induction bias could impact plant response to various stresses (Liu et al., 2015; Powell et al., 2017). To determine the extent of induction bias (from A, B and D sub-genomes) in wheat during Fe starvation, homoeolog specific expression analysis was performed. Starting from a list of 8473 gene triplets present/expressed selected as ‘accepted triplets’ (Supplementary Table S4), most of homoeolog triplets (8349) showed no significant biasness in expression (A = B = D). Out of these, 8321 triplets appeared to be unaffected by – Fe stress, while 22 and 6 homoeologous triplets were up- and down-regulated respectively. Homoeolog expression bias was observed in 124 homoeolog triplets. Eighty-seven triplets had only one of the homoeologs differentially expressed (up- or down-regulated). Sub-genome specific contribution towards this biasness within triplets is depicted in left panel of Fig. 3B. These include forty-seven in the category ‘1UP’ with A > B = D, B > A = D and D > A = B, 40 in category named ‘1DOWN’ having A < B = D, B < A = D or D < A = B. Table 2 provides the list of genes with significant induction predominance occurring from the A and B genomes in response to Fe starvation. Few of the prominent transcripts exclusively induced by these two genomes include transcripts related to MYB TFs, metal transporters, zinc transporters, RINGH-H2 type proteins, genes belonging to major facilitator superfamily proteins etc. Additionally, 37 triplets had two of the homoeologs differentially expressed while the third showed normal expression even under Fe stress (Fig. 3B; right panel). ‘2UP’ category includes AB > D, AD > B and BD > A while ‘2DOWN’ includes AB < D, AD < B and BD < A (Fig. 3B). This suggests that during Fe starvation, the additive homoeolog contribution from either A or B sub-genome was the highest.

**Table 2:** List of genes showing genome induction bias contribution from either A or B sub-genome. Expression changes within triads w.r.t. iron stress were observed to identify the presence of differential regulation among homoeologs from each triad, thus contributing towards genome induction bias. Table gives descriptive list of homoeologs that were up/down-regulated under iron starvation only in the A sub-genome, followed by those only in the B sub-genome.

| Gene | logFold change | RAP-DB description |
| --- | --- | --- |
| **A genome** |  |  |
| TRIAE_CS42_3AL_TGACv1_197291_AA0665720 | 3.01 | Myb transcription factor domain containing protein. |
| TRIAE_CS42_2AL_TGACv1_094922_AA0304900 | 2.8 | Similar to Prolyl endopeptidase (EC 3.4.21.26) (Post-proline cleaving enzyme) (PE). |
| TRIAE_CS42_5AL_TGACv1_379416_AA1256390 | 2.44 | Similar to Solute carrier family 35, member F1. |
| TRIAE_CS42_2AL_TGACv1_094606_AA0300340 | 2.18 | RmlC-like jelly roll fold domain containing protein. |
| TRIAE_CS42_3AS_TGACv1_211026_AA0683370 | 3.36 | Protein of unknown function DUF1399 family protein. |
| TRIAE_CS42_7AL_TGACv1_557470_AA1781720 | 3.47 | Heavy metal-transporting PIB-ATPase, Root-to-shoot cadmium (Cd) translocation |
| TRIAE_CS42_6AL_TGACv1_471682_AA1512850 | 2.26 | Similar to CRT/DRE binding factor 1. |
| TRIAE_CS42_6AS_TGACv1_485332_AA1543320 | 1.85 | Zinc/iron permease family protein. |
| TRIAE_CS42_2AL_TGACv1_092944_AA0267670 | 2.43 | Heavy metal transport/detoxification protein domain containing protein. |
| TRIAE_CS42_5AL_TGACv1_375378_AA1220660 | 2.21 | Similar to SUSIBA2 (WRKY protein). |
| TRIAE_CS42_1AL_TGACv1_000708_AA0017440 | 2.57 | Chlorophyll a-b binding protein 2, chloroplast precursor (LHCII type I CAB-2) (LHCP). |
| TRIAE_CS42_2AL_TGACv1_093154_AA0273510 | 1.86 | C2 domain containing protein. |
| TRIAE_CS42_5AL_TGACv1_374025_AA1188170 | 2.66 | Nodulin-like domain containing protein. |
| TRIAE_CS42_3AL_TGACv1_197522_AA0666570 | 2.22 | Lipase, GDSL domain containing protein. |
| TRIAE_CS42_7AL_TGACv1_556100_AA1755640 | 2.34 | Bifunctional inhibitor/plant lipid transfer protein/seed storage domain containing protein. |
| TRIAE_CS42_4AS_TGACv1_306527_AA1009640 | 2.2 | Multi antimicrobial extrusion protein MatE family protein. |
| TRIAE_CS42_7AS_TGACv1_570336_AA1834060 | 1.98 | Major facilitator superfamily protein. |
| TRIAE_CS42_2AL_TGACv1_093456_AA0280470 | 2.15 | Glycolipid transfer protein domain domain containing protein. |
| TRIAE_CS42_1AL_TGACv1_002205_AA0039730 | -2.09 | Similar to IAA8 (Fragment). |
| TRIAE_CS42_4AL_TGACv1_290815_AA0989640 | -2 | Lipase, class 3 family protein. |
| TRIAE_CS42_5AL_TGACv1_374561_AA1203290 | -2.24 | Cinnamyl alcohol dehydrogenase (EC 1.1.1.195). |
| TRIAE_CS42_5AL_TGACv1_374359_AA1197930 | -2.36 | Similar to Lipoygenase L-2 (EC 1.13.11.12). |
| TRIAE_CS42_6AL_TGACv1_472625_AA1524260 | -1.89 | RAG1-activating protein 1 homologue domain containing protein. |
| TRIAE_CS42_7AL_TGACv1_558250_AA1791610 | -2.32 | Similar to Pleiotropic drug resistance protein 3. |
| TRIAE_CS42_2AL_TGACv1_097246_AA0323500 | -1.72 | Similar to Peroxidase (EC 1.11.1.7). |
| TRIAE_CS42_7AL_TGACv1_556210_AA1758330 | -2.21 | Similar to Kaurene synthase A (Fragment). |
| TRIAE_CS42_1AL_TGACv1_000555_AA0014640 | -1.84 | No apical meristem (NAM) protein domain containing protein. |
| TRIAE_CS42_6AL_TGACv1_471077_AA1502240 | -1.79 | Similar to OSIGBa0145M07.8 protein. |
| TRIAE_CS42_7AL_TGACv1_559906_AA1801190 | -1.68 | Similar to H0801D08.12 protein. |
| TRIAE_CS42_7AL_TGACv1_558101_AA1790150 | -1.76 | Similar to Acyl-ACP thioesterase (Fragment). |
| TRIAE_CS42_6AL_TGACv1_472321_AA1520860 | -2.06 | Similar to Subtilisin-like protease (Fragment). |
| **B Genome** |  |  |
| TRIAE_CS42_5BS_TGACv1_423346_AA1374840 | 2.19 | Similar to Calmodulin NtCaM13. |
| TRIAE_CS42_3B_TGACv1_224030_AA0791180 | 2.2 | Similar to IN2-2 protein. |
| TRIAE_CS42_2BL_TGACv1_129296_AA0377070 | 2.46 | Similar to OSIGBa0127A14.7 protein. |
| TRIAE_CS42_6BL_TGACv1_499646_AA1588130 | 2.32 | TGF-beta receptor, type I/II extracellular region family protein. |
| TRIAE_CS42_7BS_TGACv1_592587_AA1940840 | 2.25 | Similar to RING-H2 finger protein ATL1R (RING-H2 finger protein ATL8). |
| TRIAE_CS42_2BS_TGACv1_146290_AA0461590 | 2.64 | NA |
| TRIAE_CS42_5BL_TGACv1_404610_AA1306090 | 1.84 | Similar to Senescence-associated protein SAG102. |
| TRIAE_CS42_5BL_TGACv1_405319_AA1324840 | 2.77 | Similar to Transporter associated with antigen processing-like protein. |
| TRIAE_CS42_7BL_TGACv1_577614_AA1879540 | 2.69 | Peptidase A1 domain containing protein. |
| TRIAE_CS42_4BS_TGACv1_329166_AA1098520 | 2.32 | Similar to Alcohol dehydrogenase. |
| TRIAE_CS42_4BL_TGACv1_321683_AA1064050 | 2.56 | Protein of unknown function DUF1262 family protein. |
| TRIAE_CS42_7BL_TGACv1_591489_AA1920550 | NA | Similar to zinc transporter 4. |
| TRIAE_CS42_7BL_TGACv1_577301_AA1871590 | 2.41 | Delta-tonoplast intrinsic protein. |
| TRIAE_CS42_4BS_TGACv1_329309_AA1100040 | NA | Similar to Major facilitator superfamily antiporter. |
| TRIAE_CS42_2BL_TGACv1_129348_AA0379680 | 2 | Hypothetical conserved gene. |
| TRIAE_CS42_2BS_TGACv1_148847_AA0495340 | 4.34 | Helix-loop-helix DNA-binding domain containing protein. |
| TRIAE_CS42_2BL_TGACv1_130820_AA0418390 | 1.88 | NA |
| TRIAE_CS42_1BL_TGACv1_031794_AA0120680 | -2.79 | Divalent ion symporter domain containing protein. |
| TRIAE_CS42_5BL_TGACv1_406838_AA1350360 | -2.29 | Similar to anther-specific proline-rich protein APG. |
| TRIAE_CS42_2BL_TGACv1_130182_AA0405730 | -2.09 | Protein of unknown function DUF3741 domain containing protein. |
| TRIAE_CS42_4BS_TGACv1_330685_AA1108490 | -2.83 | Serine/threonine protein kinase-related domain containing protein. |
| TRIAE_CS42_2BS_TGACv1_147909_AA0488910 | -2.28 | Delayed-early response protein/equilibrative nucleoside transporter family protein. |
| TRIAE_CS42_3B_TGACv1_224656_AA0799480 | -1.83 | Glyoxalase/bleomycin resistance protein/dioxygenase domain containing protein. |
| TRIAE_CS42_7BL_TGACv1_577476_AA1876230 | -2.93 | Protein kinase, catalytic domain domain containing protein. |
| TRIAE_CS42_6BL_TGACv1_499809_AA1592210 | -1.88 | Cupredoxin domain containing protein. |
| TRIAE_CS42_7BL_TGACv1_576755_AA1853830 | -2.04 | Cellulase (EC 3.2.1.4). |
| TRIAE_CS42_2BL_TGACv1_130686_AA0416110 | -2.3 | Similar to OSIGBa0096P03.3 protein. |
| TRIAE_CS42_2BS_TGACv1_146035_AA0453750 | NA | Conserved hypothetical protein. |
| TRIAE_CS42_1BL_TGACv1_030699_AA0098220 | -1.91 | Lipase, class 3 family protein. |
| TRIAE_CS42_2BL_TGACv1_129634_AA0391150 | -1.78 | 2OG-Fe(II) oxygenase domain containing protein. |

**Fig. 3.**
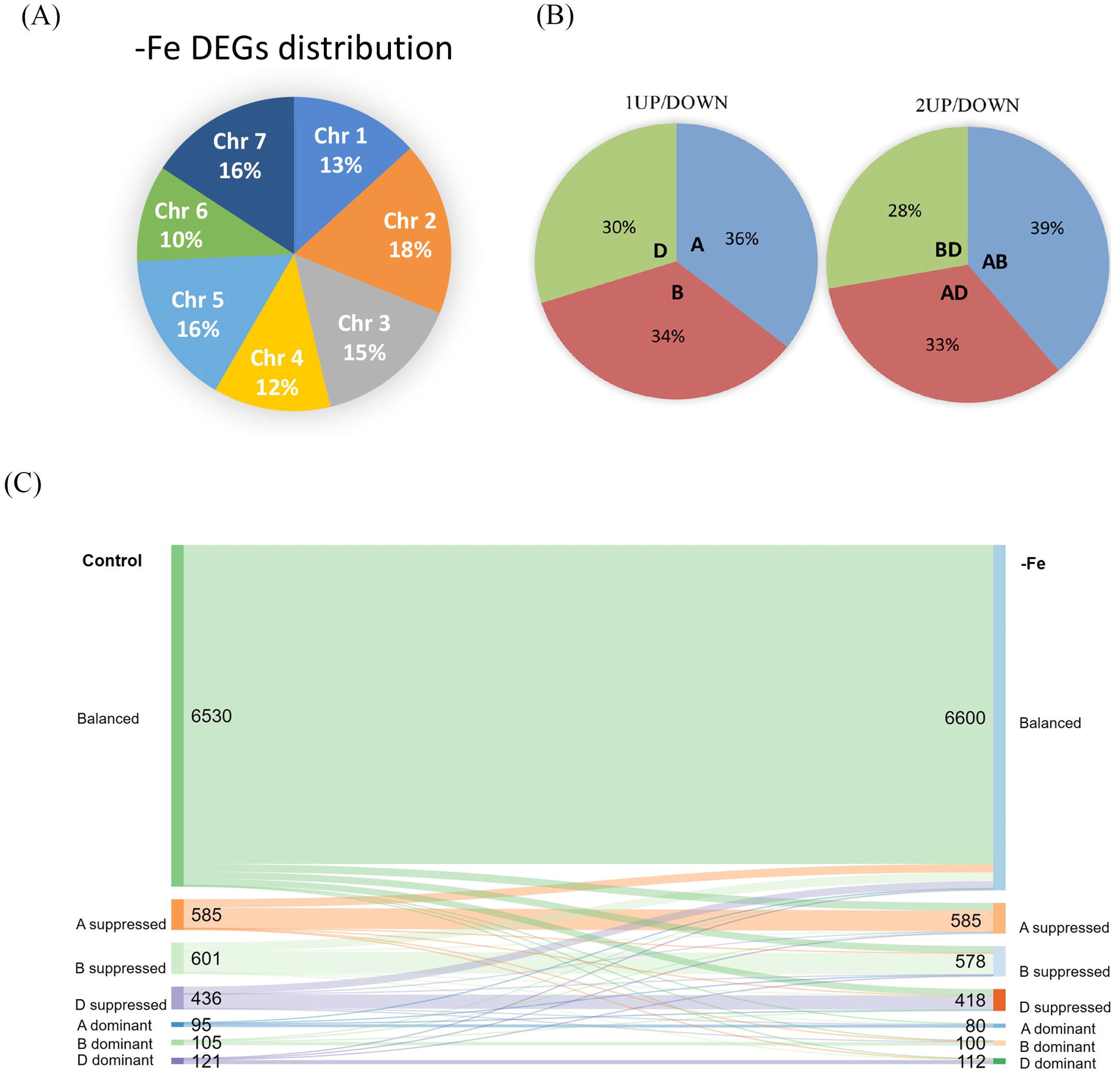
Genomic distribution and homoeolog bias studies during Fe starvation. (A) Chromosomal inclination of DEGs. (B) Pie charts showing (left panel) sub-genomic distribution of genome induction bias in triads where one of the homoeologs was up/down-regulated. A, B and D depict the sub-genome to which the DE homoeolog belongs; (right Panel) distribution of triads for which two of the homoeologs were differentially expressed and the third one had normal expression. AB refers to the triads for which up/down regulation was observed in the homoeologs belonging to A and B sub-genomes, while the D sub-genome homoeolog behaved normal w.r.t control. (C) Sankey diagram depicting the homoeolog expression bias in Control condition and Fe starvation. Homoeolog triads were classified into the defined seven categories based on relative normalised expression within each triad. Nodes flowing from Control to –Fe (Fe starvation) represent the triads with same (flow to same category) as well as changed (flow to a different category) expression patterns across both conditions. Distinct colors represent the flow of triads belonging to the seven categories form control condition into same category under –Fe or transition into a different category under –Fe.

For polyploid genomes such as wheat, the interaction of its sub-genomes is known to affect the final phenotype or contribute towards trait development (Borrill et al., 2015). To check the homoeolog/sub-genome expression biasness under *–*Fe condition, effect on the expression of transcripts from the genomes was performed by comparing the relative normalised expression for each homoeolog within a triad. This resulted seven combinations including one balanced category and six homoeolog specific dominance or suppression categories. Our analysis reflected that most of the triads were falling under the balanced category (Table 3 and Fig.3C). This category was represented by 77% and 77.89% of the total triads for control and Fe starvation condition. Triads with unbalanced expression varied in the range of 0.94 to 7.09 for all the 6 sub-categories across the two conditions (Table 3 and Fig.3C). Our analysis revealed that maximum genome specific expression biasness was observed for A and B on both the conditions. Interestingly, D genome was least suppressed with a representation of 5.15% of the total triads taken in consideration in control when compared to 4.93% in Fe starvation. 89% triads showed conserved balanced/unbalanced contribution across both conditions. Overall, a higher relative abundance of D genome (Control: 33.94%; –Fe: 33.95%) was noted as compared to the A (Control 33.06%; –Fe: 33.00%) and B (Control: 33.04%; –Fe: 33.95%) genomes.

**Table 3:**
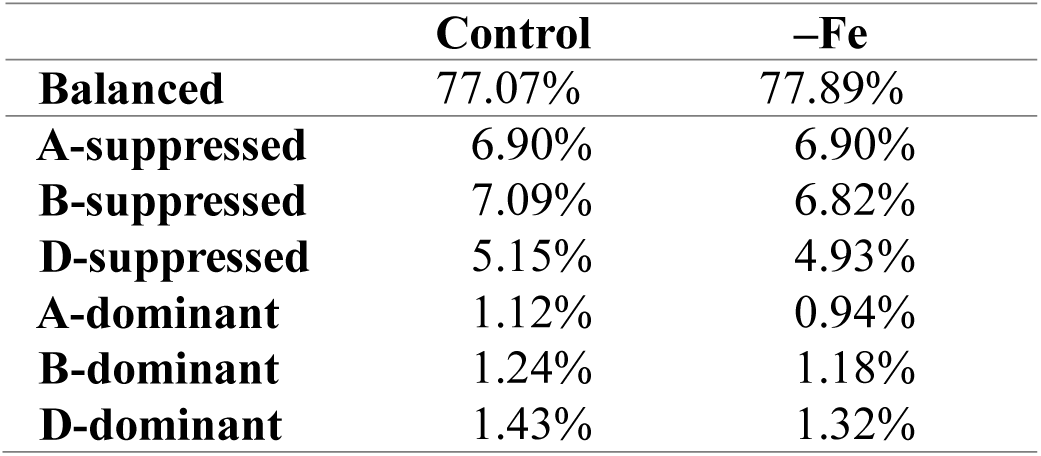
Percentage of homoeolog triads categorised into ideal genome expression bias categories in control and Fe starved conditions.

### Identification of DEGs pinpoints the prolific expression of genes involved in Strategy-II mode of Fe uptake

To identify the transcripts differentially expressed in roots in response to Fe starvation, top 50 genes either up-regulated or down-regulated were shortlisted (Fig. 4). Expression of most of the highly up-regulated transcripts ranged from 12 to 4.8 log2 fold change (Supplementary Table S3) indicating their higher fold accumulation under Fe starvation compared to the control conditions. As observed in the top 50 upregulated genes, the predominance of pathway genes encoding the components for Strategy-II mode Fe uptake was observed across all the 3478 upregulated genes. Categorically, these highly induced genes are known to be involved during Strategy-II mode of iron uptake, including the sub-family of nicotinamine synthase (NAS) and deoxymugineic acids (DMA) biosynthesis genes (Fig. 4, left panel). Other important genes include genes belonging to the Major Facilitator Superfamily including an ABC transporter, zinc-induced facilitator like transporters (ZIFL), sulphate transporters. Therefore, under Fe starvation, the DMAS biosynthesis genes were highly induced. Similarly, YSL genes were significantly induced under Fe starvation condition. All the homoeologs of *TaYSL9* and *TaYSL1A* showed especially high expression under starvation condition (Supplementary Table S5). On the similar lines, genes encoding for NRAMP also showed high transcript abundance in roots subjected to Fe starvation (Supplementary Table S5). The validation of the expression of few Strategy-II uptake genes was also done by quantitative real time-PCR (qRT-PCR). Our results for qRT-PCR validate our inference from the RNAseq analysis. Almost all of the Strategy-II genes tested for their expression showed a very high fold expression in Fe starved roots as compared to the control (Supplementary Fig. S1). Highest expression was obtained for ZIFL4, DMAS1, NAAT1, NAS1 those are the prime components for Strategy-II mediated uptake. Other transcripts encoding for thaumatin like proteins etc. were also induced under Fe limiting condition.

**Fig. 4.**
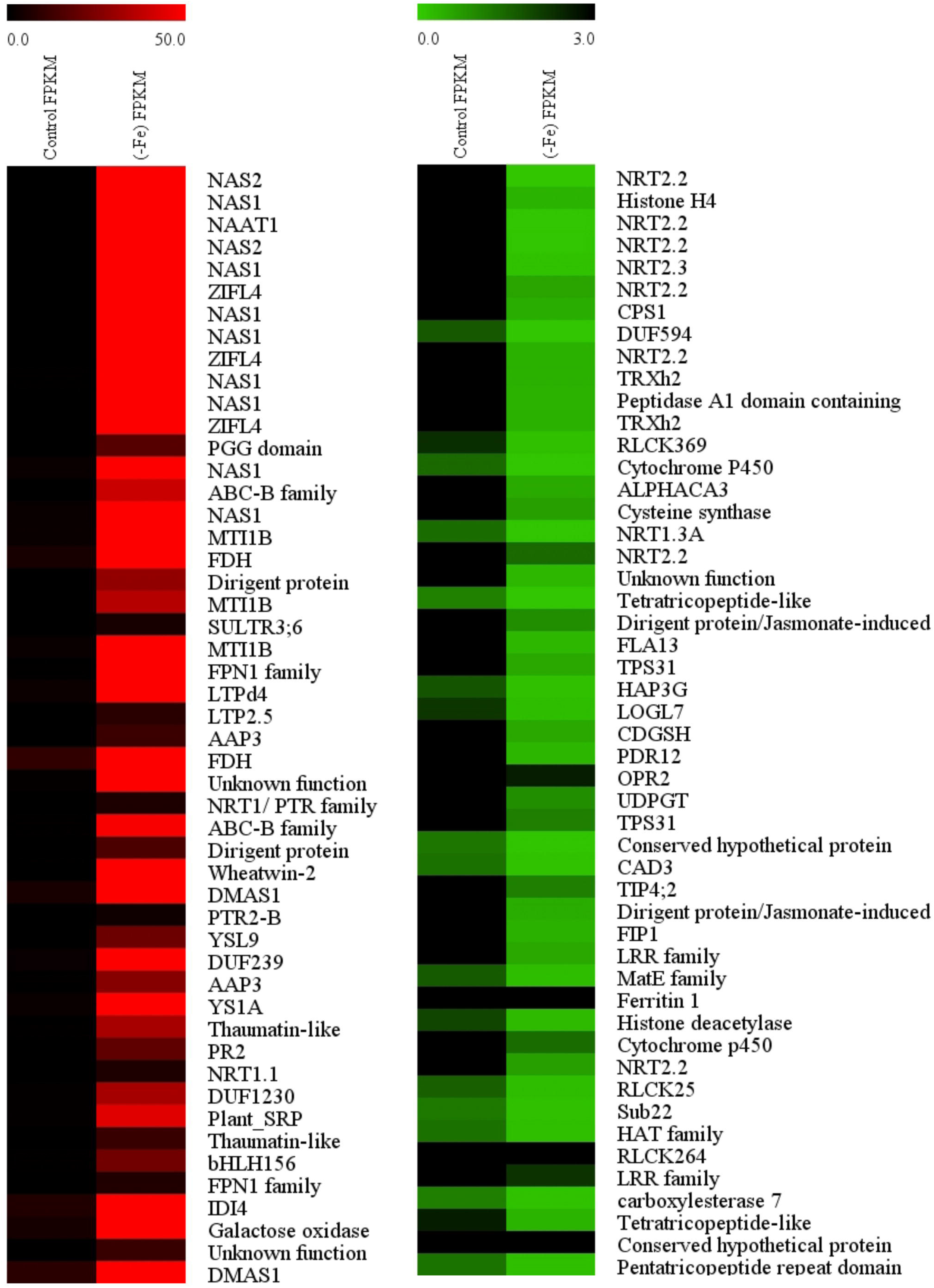
Top up-regulated and down-regulated genes in –Fe condition, annotated via KOBAS using rice RAP-DB/RefSeq annotations as reference. The heat map shows top 50 genes those are highly up-regulated (red-left panel) and down-regulated (green-right panel) identified in wheat roots under –Fe condition with respect to control. For expression analysis, FPKM values were obtained using Cufflinks, and CuffDiff was used to identify DEGs by calculating significant changes in transcript expression between the stressed and normal samples (FDR≤0.05).

Transcripts of genes involved in the Strategy-I mode of Fe uptake were also analysed (Supplementary Table S6). An important component of Strategy-I pathway that includes H^+^-ATPase (AHA) subfamily genes was not differentially expressed under Fe limiting condition (Supplementary Table S6). This was in agreement with the qRT-PCR analysis where, either no change or down-regulation of AHA genes was observed (Supplementary Fig. S1) in wheat roots. Metallo-reductases are important components of Strategy-I represented by ferric-chelate reductase (FRO). Most of the transcripts encoding for wheat FROs do not show significant changes in starved roots as compared to control except for one transcript. Interestingly, iron regulated transporters (IRT; TRIAE_CS42_7DS_TGACv1_622068_AA2032200; TRIAE_CS42_4AL_TGACv1_289466_AA0971640) were significantly expressed in Fe starved wheat roots, suggesting their involvement in wheat under metal stress. The high expression of *TaIRT1*was also confirmed by qRT-PCR (Supplementary Fig. S1). This suggest that IRT function might be conserved among the plant species.

In addition to this, forty-five genes showed exclusive transcript abundance under Fe starvation (Supplementary Table S3). No transcript for any of these genes was detected in the control root samples, suggesting their high specificity for Fe response. Some of the transcripts responding to Fe starvation encode for metallothionine, metal transporters, vacuolar iron transporters (VIT1) and nicotianamine synthase 2 (NAS2). This suggests that in wheat few components involved in Fe uptake could actually respond exclusive to Fe starvation. The fold expression levels of genes those were highly down-regulated ranged from −7.1 to −2.85-fold at the level of log2 scale (Supplementary Table S3). Interestingly, multiple genes encoding for nitrate transporters were highly down-regulated under Fe limiting condition (Fig. 4, right panel). Two genes encoding for cytochrome-P450 also showed down-regulation. Among the others, genes encoding for Histone deacetylase, peptidases A1 containing domain protein, cinnamyl alcohol dehydrogenase, dirigent protein show significant down-regulation.

### Functional enrichment network of Fe starvation related genes

Gene ontology annotations and classification of DEGs was performed to get the overview of processes those are representation of cellular, molecular and biological functions. Analysis was further extended to cluster analysis using Cytoscape plugins, BINGO and Enrichment Map (Maere et al., 2005; Merico et al., 2010). A total of 5854 DEGs were found to be differentially-expressed in response to Fe starvation; significant GO categories were assigned to all DEGs. The DEGs annotated for GO terms were visualized using WEGO tool (Fig. 5A, Supplementary Table S7). Most enriched GO terms in the “cellular component” category were membrane and intracellular organelle. Other overrepresented terms included catalytic activity and ion as well as organic compound binding for the “molecular function” category, while metabolic processes related to nitrogen as well as other cellular processes were observed for “biological process” category (Fig. 5A). Overall, catalytic activity and binding activities were most significantly enriched GO terms in –Fe condition (Fig. 5B). Further mapping of upregulated genes to databases such as, Kyoto encyclopedia of genes and genomes (KEGG) pathway (Xie et al., 2011) and MapMan (Thimm et al., 2004) revealed enrichment related to phenyl-propanoid biosynthesis, amino acid biosynthesis and carbon metabolism, and glutathione metabolism (Fig. 5C and Supplementary Fig. S2). The role of glutathione in response to Fe starvation is intriguing and deserves further investigations.

**Fig. 5.**
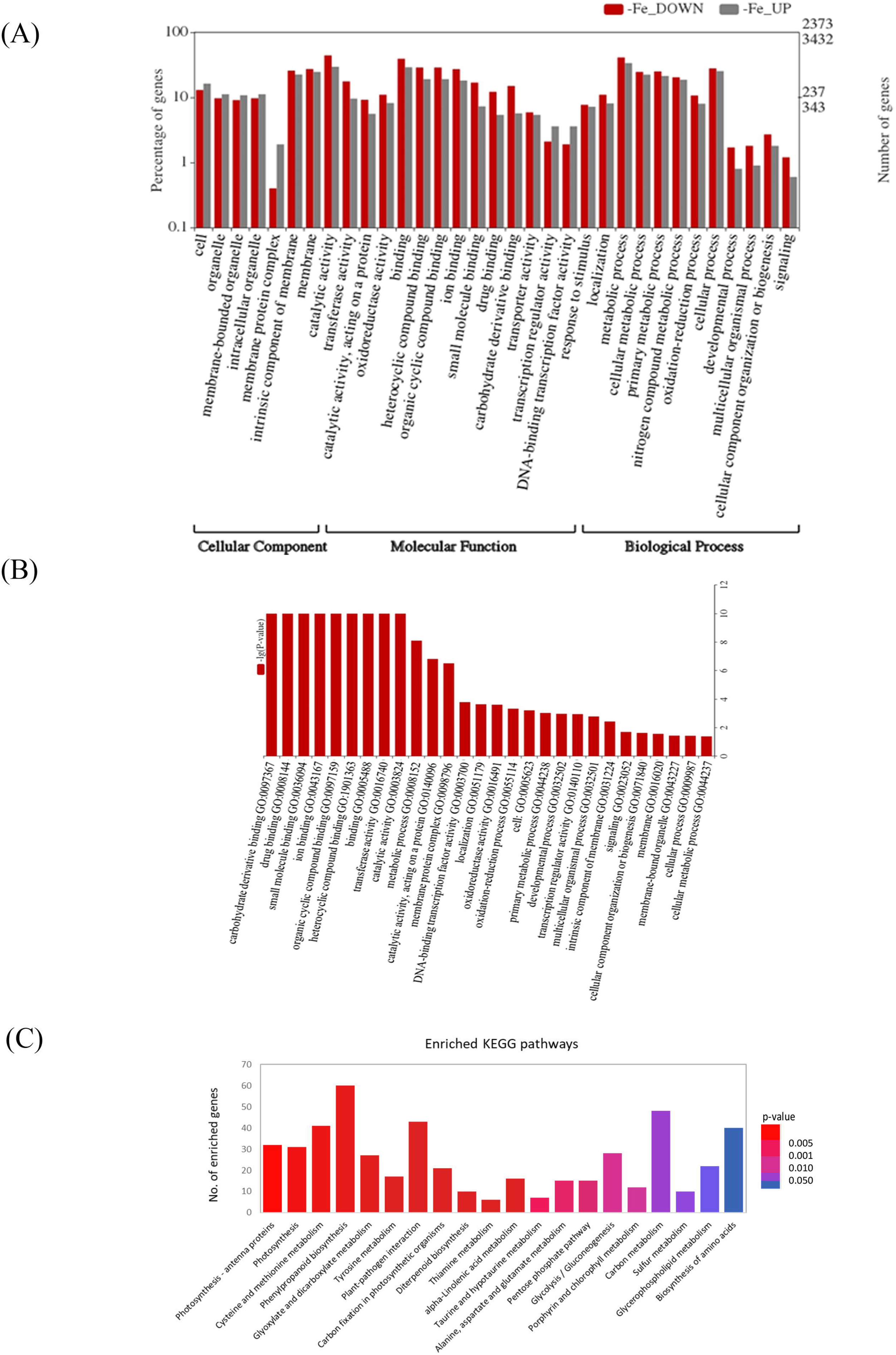
Gene ontology (GO) categorization of the differentially expressed genes and its analysis. (A) WEGO plot describing GO annotation and classification of DEGs, with y-axis showing the percentage of genes belonging to respective GO terms (grey bars for down-regulated genes and red bars for up-regulated genes) as well as the right plane ticks for y-axis depicting the number of both up-and down-regulated genes represented by the respective percentages on the left plane. Percentage and number of genes were calculated for the three broad main categories listed on the x-axis; (B) Enriched GO terms in DEGs under –Fe condition, y-axis depicting the significance of GO term enrichment. (C) Figure showing top 20 enriched KEGG pathways represented by the up-regulated genes under iron starvation condition. x-axis showing names of the pathways, y-axis representing the number of genes enriched in respective pathways.

GO category “integral to membrane” was found to be associated with pathways clustered into “metal ion and trans-membrane transport” and “photosynthesis” related GO categories (Fig. 6A). Clustered pathways involved in “response to nutrient levels” and “Sulphur amino acid metabolism” was also enriched (Fig. 6A). A few other up-regulated clustered pathway genes involved fall into the category “response to phosphate starvation”. In the enriched network of down-regulated DEGs, genes involved in iron and other metal homeostasis were associated with two different clusters, mainly included genes related to lipid, ketone and carboxylic metabolism, in addition to those involved in nitric acid and salicylic acid response (Fig. 6B). A second cluster contains transcripts related to purine and adenine nucleotide binding and pyrophosphatase activity. Thus, whole set of significant up-regulated and down-regulated DEGs were clustered in distinct cellular processes suggesting differential transcriptional response under Fe starvation.

**Fig. 6.**
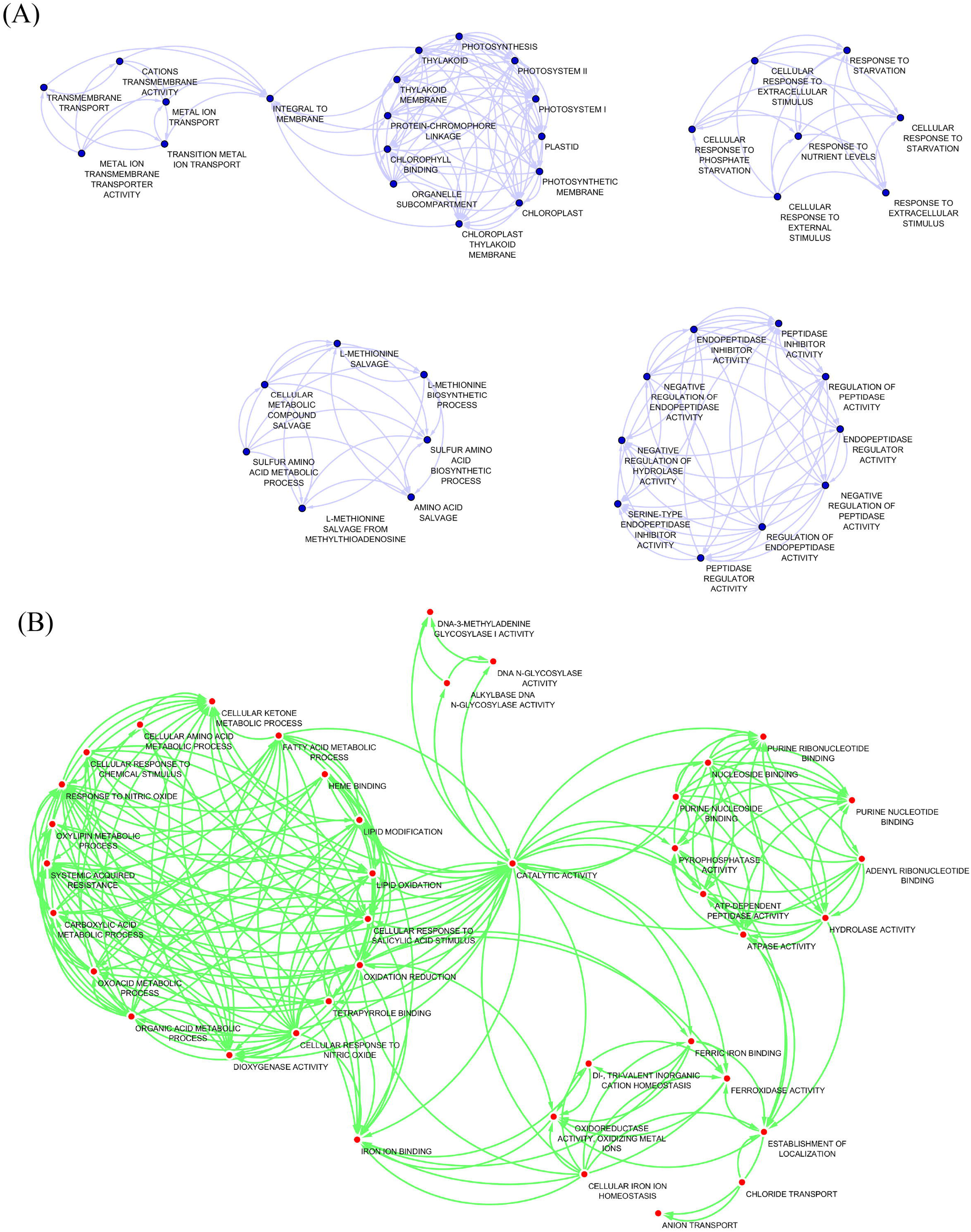
Co-expression/hub genes and function enrichment network for identified DEGs. Function enrichment network for DEGs associated under Iron starvation in wheat roots with high significance (FDR ≤ 0.05) for (A) Up-regulated and (B) Down-regulated genes. Enriched GO functional categories are clustered with correlated DEGs and represented by node circles.

### Identification of transcriptional regulatory genes during Fe starvation

To address an important question on how the –Fe signal is connected to the transcriptional machinery in wheat, genes encoding TFs were identified. Earlier the TFs involved in the response to Fe starvation were identified in the model plant Arabidopsis and rice including POPEYE (PYE), basic helix-loop helix (bHLH), ethylene insensitive-3 (EIN3), EIN3-like1 (EIL1) (Long et al., 2010, Mai et al., 2016, Bauer et al., 2011, Ivanov et al., 2012; Ogo et al., 2011; Bashir et al., 2014). Consistently, our transcriptome analysis showed that all homoeologs for PYE encoding a bHLH transcription factor and BRUTUS-like (BTS) known to encode a putative hemerythrin E3 ligase protein in Arabidopsis and rice were highly expressed in wheat roots under Fe starvation (Supplementary Table S8). Analysis of these category of genes led to identification of 41 significantly up-regulated TF family members (>log2FC) (Supplementary Table S8). The TF family members belong to categories like APETALA2 ethylene-responsive element binding proteins (AP2/EREBP), WRKY, C2H2, Zinc finger proteins (including C3HC4), NAM, bHLH hemerythrin and U-box (Fig. 7A). Out of these, transcripts encoding for AP2/EREBP followed by WRKY and bHLH were the most abundant in wheat. Genes encoding for AP2-EREBP, WRKY and C2H2 type of TFs also showed high predominance (Supplementary Table S8 and Supplementary Fig. S3). However, genes encoding for TFs such as MADS, PHD and homeobox (HB) domain containing proteins were highly represented in the list of down-regulated transcripts. Finally, to ascertain regulatory functions under iron starvation, enrichment mapping was performed for the significantly expressed DEGs those were classified into transcriptional factors. Interestingly, genes involved in regulation of gene expression were clustered with primary and nucleotide metabolism related genes in both positive and negative correlation. This functional cluster was in positive correlation with TFs involved in gene ontology (GO) categories such as “vacuolar transport” and “DNA topoisomerase III activity”. However, two separate clusters “cellular response to auxin stimulus and signalling” and “cell-fate specification” were in negative correlation with “regulation of gene expression” and “primary and nucleotide metabolism” (Fig. 7B). This suggested an important regulatory role of TFs involved in auxin signalling and cell-fate specification in Fe starvation response control, thereby modulating the network of Fe homeostasis.

**Fig. 7.**
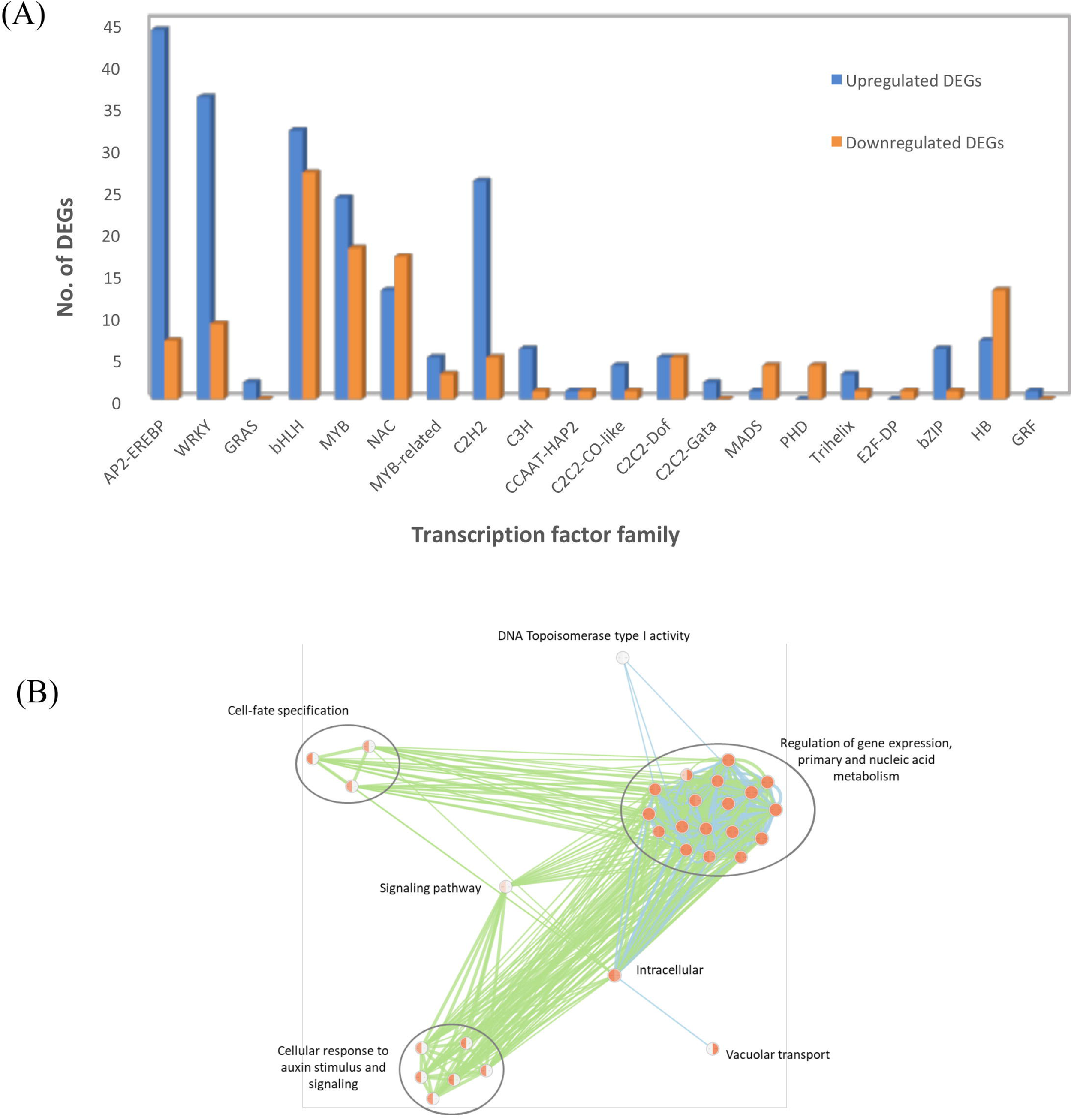
Transcriptional factors (TFs) significantly associated with Fe starvation (FDR ≤ 0.05) in wheat roots. (A) List of TFs differentially expressed in response to Fe starvation stress. Blue bars represent up-regulated and red bars represent down-regulated TFs. (B) Co-expression/hub genes and network analysis using Fe responsive TFs (FDR ≤ 0.05) in wheat roots. Circles indicate the processes associated with it (green for down-regulated genes; Blue for up-regulated genes). Enriched GO functional categories are clustered with correlated TFs and represented by node circles.

### Glutathione-S-transferases are involved in the response to Fe starvation in wheat

Glutathione mediated conjugation of multiple metabolites plays important role during the metal stress (Zhang et al., 2013). Our transcriptome data revealed the enhanced expression of multiple glutathione-S-transferases (GST) in Fe-starved wheat root as compared to control (Supplementary Table S9). To correlate the expression response with its enzymatic activity, temporal response of GST was measured in wheat roots under –Fe conditions. Using the GST functional assay, the activity was determined in wheat roots of plants grown for 10, 15 and 20 days under Fe starvation. Our activity assays showed significant increase in the GST activity under –Fe condition, which peaked at 10 and 20 days after the beginning of Fe starvation compared to control plants (Fig. 8A). Therefore, our experiments validate the increased GST transcript abundance with enhanced glutathione activity. These results indicate an important role of glutathione in response to Fe starvation in wheat roots.

**Fig. 8.**
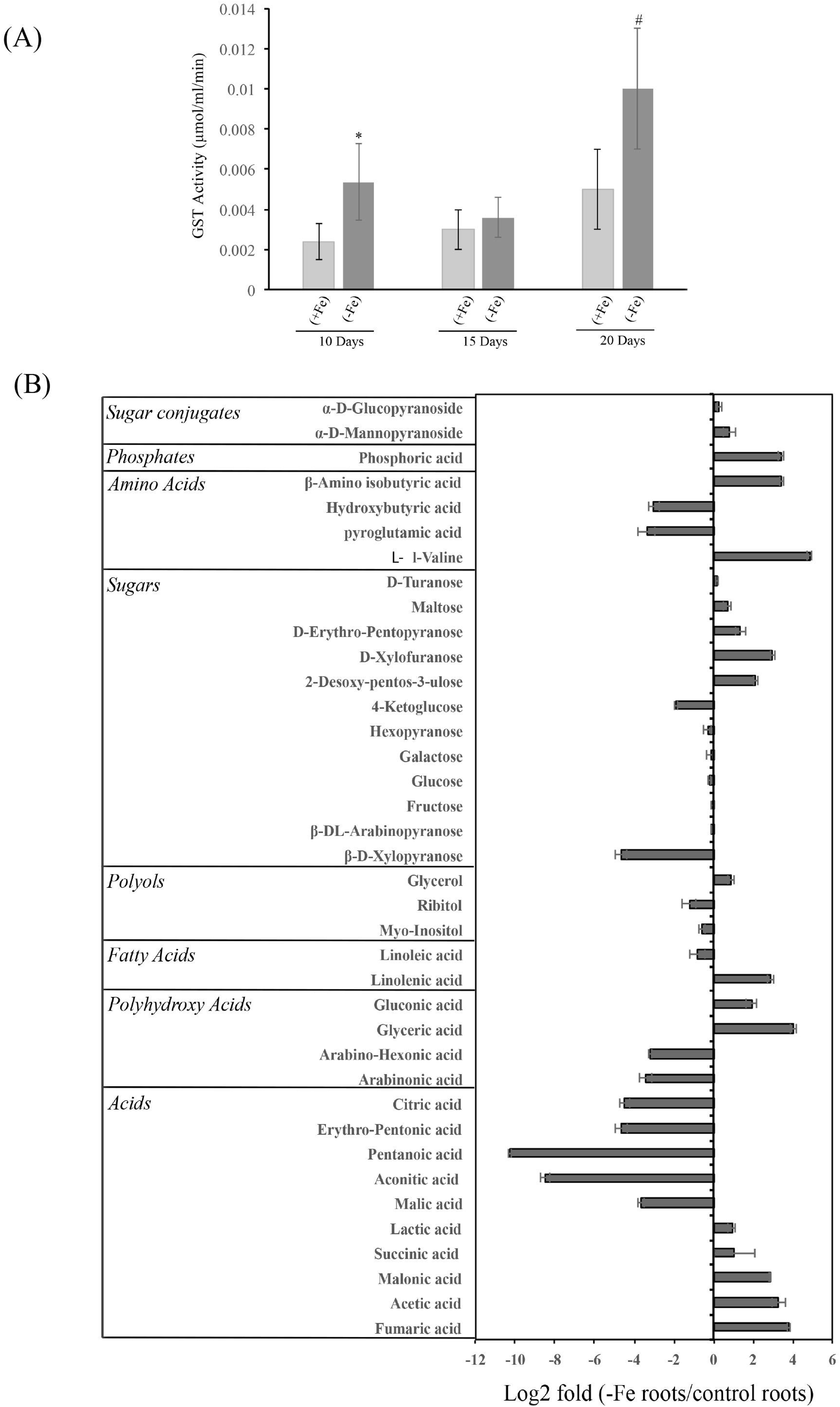
Measurement of GST activity and metabolite profiling of wheat roots subjected to Fe starvation. (A) Glutathione-S-transferase activity of wheat roots under Fe starvation (–Fe) and control (+Fe) condition. (B) Metabolite profiling of amino acids, sugars, polyols, organic acids and related compounds. Change in abundance of significant (P < 0.05) metabolites identified by GC-MS in Fe-starved roots. Abundance variation of each metabolite is represented in Log2 fold values of response ratio (–Fe /+Fe) of metabolite concentrations. Values are means of three biological replicates with bar representing Log ratio of standard error. * indicate significant difference at p<0.01; # indicate significant difference at p<0.05.

### Fe starvation causes an accumulation of organic acids and polyhydroxy acids in wheat

To obtain a comparative insight of the metabolite profile of wheat roots of plants grown in absence or presence or Fe; GC-MS profiling analysis was performed. Metabolites were extracted from the roots of three replicate pools of plants each containing eight seedlings. Qualitative processing of each chromatogram for peak area and identification was performed in MassHunter version B.05.00 software coupled with NIST11 compound library. The compound annotation was determined by comparing individual resolved peaks to library searches based on mass spectra and compounds chromatographic retention indices. Interestingly, analysis resulted in the identification of 54 metabolites and further 39 annotated metabolites were analysed for their response ratio (Supplementary Table S10). To compare the change under Fe-starvation, metabolite abundances (–Fe roots/control roots) were calculated and expressed in Log2 fold change values. Fe-starvation significantly affected accumulation of 22 metabolites that includes organic acids, polyhydroxy acids, amino acids and some of the sugars, fatty acids and phosphates (Fig. 8B). Amongst treatment-specific changes, few organic acids such as fumaric acid, acetic acid and malonic acid showed higher level of accumulation in Fe starved roots as compared to the control. In contrast, citric acid, malic acid, valeric acid and aconitic acid were significantly lowered in Fe starved roots when compared to control samples. In Fe-starved roots, the accumulation of amino acids mainly, L-Valine showed significant increase while hydroxyl butyric acid and pyroglutamic acid was lowered in comparison with control roots. Polyhydroxy acids like gluconic acid and glyceric acid were also significantly high in Fe starved roots, whereas level of hexonic acid and arabinoic acid was found to be low. Taken together, our results showed that during Fe starvation wheat roots undergo reprograming for metabolic changes to maintain the Fe-flux.

## Discussion

How plants maintain nutrient homeostasis is a fascinating question in plant biology. In this direction, *A. thaliana*, with its fully sequenced small genome has provided some basic information. Developing our knowledge on nutrient homeostasis in crops, especially those having complex genomes, including hexaploid wheat is more challenging. Recently, the release of genome sequence, generation of transcriptome data and the advancement of different functional tools have provided much needed impetus in this direction (Borrill et al., 2015). The current study was undertaken to gain insight into the response of hexaploid wheat exposed to Fe starvation that is known to severely affect crop production. In this study, transcriptome of wheat (cultivar C306) in response to Fe starvation was generated. Our analysis revealed that; (a) wheat utilizes primarily Strategy-II mode of Fe uptake, (b) accumulates transcripts encoding for methionine-salvage pathway coupled with enhanced GST activity and (c) accumulates specific metabolites including fumarate, acetate, malonate and xylofuranose, to efficiently mobilize soil Fe. Interestingly, our systematic analysis of expression data revealed that transcripts show an induction biasness for A and B genomes of wheat in response to –Fe condition. Overall, this work provides the first comprehensive insight at the molecular level for wheat roots during Fe-deficiency.

Wheat plants subjected to Fe starvation show physiological defects such as a decrease in the root growth. This phenotype was consistent with the previous report showing negative impact on the root growth of wheat seedlings under Fe stress (Garnica et al., 2018). Our analysis also suggested down-regulation of few nitrate transporters for the time point studied. Homology searches indicated the identified nitrate transporters in wheat as *TaNRT2.2* and *TaNRT2.3* (Buchner and Hawkesford, 2014). These observations suggest that Fe starvation represses nitrate transporters, which in turn leads to lower accumulation of nitrate levels in the roots (Supplementary Fig. S4). Although impact of Fe starvation on the nitrate metabolism has not been studied in detail, yet similar decrease in nitrate levels were also observed in cucumber shoots subjected to –Fe condition (Borlotti et al., 2012). This observation further reinforces the existence of an interaction between macro and micronutrients in plants, which has recently gained attention (Rouached and Rhee, 2017; Bouain et al., 2019).

Due to the low availability of micronutrients in soil, plants are equipped in recruiting components that could participate in the two major strategies for Fe uptake (Kobayashi and Nishizawa, 2012). Cereals such as maize and rice predominantly utilize Strategy-II mode of uptake, unlike Arabidopsis, that employs Strategy-I mode of Fe uptake (Li et al., 2014; Zanin et al., 2017; Hell and Stephan, 2003). Nevertheless, a comprehensive study to identify the molecular players involved during Fe limiting conditions in wheat is lacking. Our RNAseq based analysis, strongly suggested an increase in the transcript abundance of genes for Strategy-II mode of Fe uptake. With the exception of conserved genes like *IRT1* (iron regulated transporter), prime genes for Strategy-I uptake mechanism like *FRO* (Ferric chelate reductase) and proton *H-ATPase* (AHA-like) were also present during starved condition but at very low abundance and no FRO activity has been reported for cereals. During this study, wheat root subjected to Fe starvation also does not show any FRO activity (data not shown). Therefore, our results reveal that wheat utilize Strategy-II mediated uptake of Fe and induces *IRT* genes that might be conserved for their function across the plant kingdom.

The series of events during Strategy-II uptake mechanism used by Gramineae involves secretion of PS that facilitate conjugation of Fe to form Fe-complexes. These Fe-PS complexes are imported in roots for remobilization to the different developing organs. As represented in Figure 9, the transport of these Fe-complexes occurs via membrane transporters via ZIFL genes encoding for TOM proteins (Nozoye et al., 2011; Nozoye et al., 2015). Multiple wheat ZIFL show high transcript accumulation and are closest homolog of OsTOM1 from rice, thereby speculating its role in Fe acquisition. Utilizing gain- and loss of function approaches OsTOM1 has been demonstrated to be a DMA effluxer for its role in enhancing mobilization and thereby improving Fe uptake (Nozoye et al., 2011). Altogether, based upon the high similarity and response under Fe starvation, it is tempting to propose wheat ZIFL (ZIFL4) as a functional transporter of DMA. Although, other *ZIFL*s were also significantly expressed but functional wheat transporter for DMA needs to be deciphered. PS secretion was also observed during our analysis for the roots of C306 exposed to Fe starvation. In this study a high PS release of ∼40-45 nmolg^−1^ root biomass (3 hrs)^−1^ was observed, reinforcing our RNAseq data that DMA biosynthesis genes and its transporters were highly upregulated. Control plants show accumulation of ∼1-2 nmolg^−1^ suggesting that wheat releases a basal level of PS to maintain the constant flux of Fe in roots. Upon careful analysis of the RNAseq data, it was observed that multiple such efflux transporters are represented in the list that are highly abundant under Fe starvation. During our study, we identified *TaDMAS1* encoding for deoxymugineic acid synthase that was highly responsive for Fe starvation (Supplementary TableS5). Earlier, *TaDMAS1* was reported as a gene that was broadly expressed across the tissue and was regulated during Fe-starved condition. These observations support the notion that TaDMAS1 has potential to enhance the seed iron storage capacity (Beasley et al., 2017). During our study, GO term enrichment was also observed for DMA biosynthesis pathway, transmembrane transporters and cellular response that reinforce importance of these functional categories. Surprisingly, genes pertaining to photosynthesis were also observed. This anomaly could be explained by the presence of active photosystem in the basal roots of our collected wheat tissue, similar to that observed in Arabidopsis due to the changes in the auxin/cytokinin ratio (Kobayashi et al., 2012). Secondly, during this study the wheat roots under Fe starvation show significant accumulation of two Golden-2 like (GLK) TF’s (∼logFC 1.54 and 1.55; Supplementary Table S3). GLK’s are known to improve phototrophic performance of roots and thereby enhancing root photosynthesis (Kobayashi et al., 2013). These reasons may account for the enhanced accumulation of transcripts related to photosynthesis along with other metabolic related genes.

**Fig. 9.**
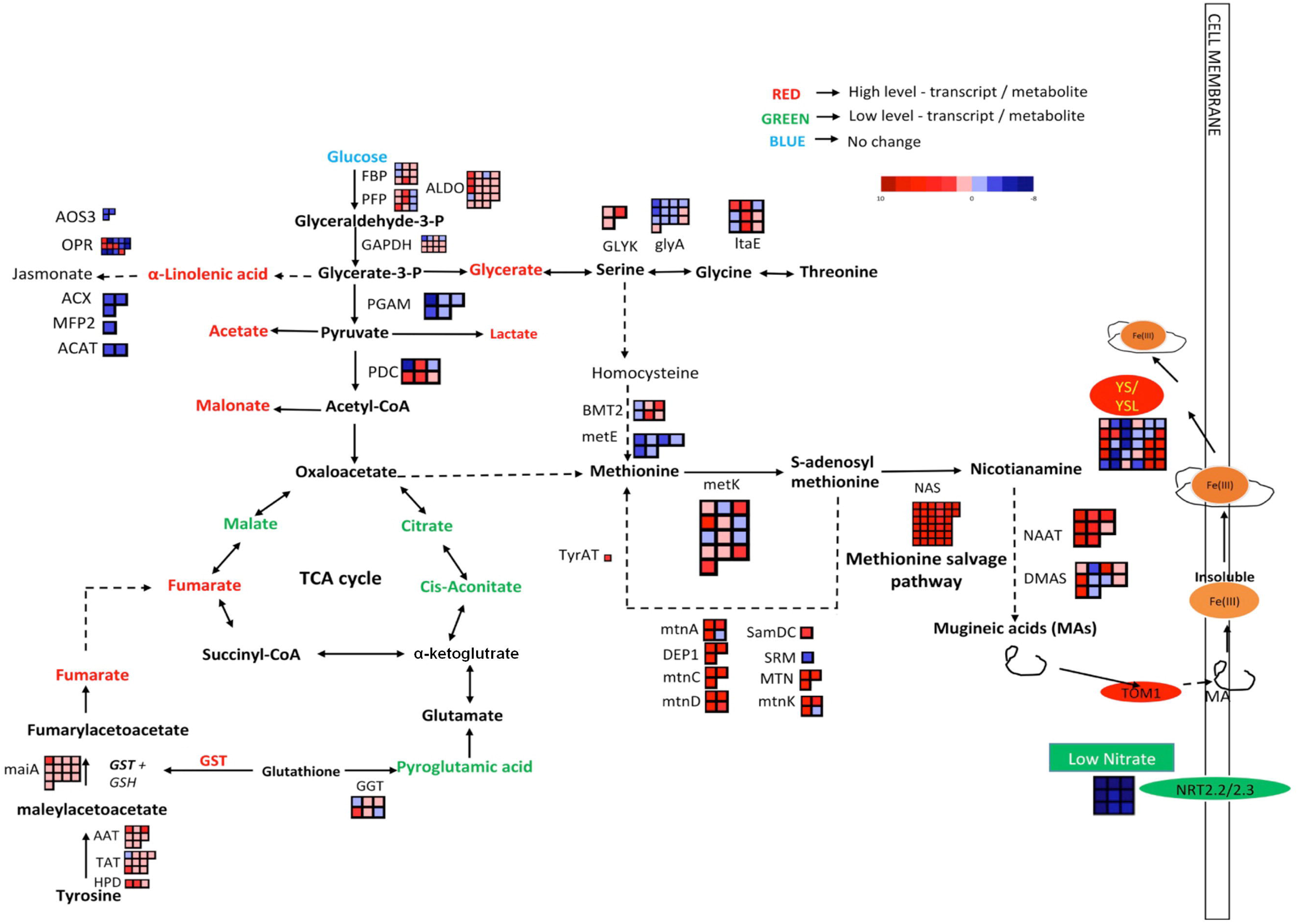
Schematic representation describing the core components involved in Fe starvation. The red font indicates the genes/metabolite those were highly up-regulated/showed high-accumulation during our study; whereas the green font indicates down-regulated/low accumulation. Methionine salvage pathway bins indicate the level of the gene expression levels for the transcript indicated next to it. Red and blue bins near pathways and steps represent the up- and down-regulation of related genes, respectively. (Abbreviation used: FBP: fructose-1,6-bisphosphatase I, PFP: diphosphate-dependent phosphofructokinase, ALDO: fructose-bisphosphate aldolase, GAPDH: glyceraldehyde 3-phosphate dehydrogenase, PGAM: 2,3-bisphosphoglycerate-dependent phosphoglycerate mutase, PDC: pyruvate decarboxylase, GLYK: D-glycerate 3-kinase, GlyA: glycine hydroxymethyltransferase, ltaE: threonine aldolase, BMT2: homocysteine S-methyltransferase, metE: homocysteine methyltransferase, metK: S-adenosylmethionine synthetase, SamDC: S-adenosylmethionine decarboxylase, SRM: spermidine synthase, MTN: 5’-methylthioadenosine nucleosidase, mtnK: 5-methylthioribose kinase, mtnA: methylthioribose-1-phosphate isomerase, DEP1: enolase-phosphatase E1, mtnC: enolase-phosphatase E1, mtnD: 1,2-dihydroxy-3-keto-5-methylthiopentene dioxygenase, TyrAT: tyrosine aminotransferase, NAS: nicotianamine synthase, NAAT: nicotianamine aminotransferase, DMAS: 3’’-deamino-3’’-oxonicotianamine reductase, YSL: Yellow Stripe Like, maiA: maleylacetoacetate isomerase, AAT: aspartate aminotransferase, TAT: tyrosine aminotransferase, HPD: 4-hydroxyphenylpyruvate dioxygenase, GGT: gamma-glutamyltranspeptidase, AOS3: hydroperoxide dehydratase, OPR: 12-oxophytodienoic acid reductase, ACX: acyl-CoA oxidase, MFP2: enoyl-CoA hydratase, ACAT: acetyl-CoA acyltransferase 1)

Wheat is a hexaploid crop with three genomes and therefore studying the homoeolog induction and expression bias could provide an insight on the regulation of the transcript under a given biotic and abiotic stress. Therefore, we subjected our RNAseq data to these analyses. In depth analysis confirms the minimal suppression for the expression bias for the D-genome derived transcripts as compared to A or B. These observations support the previous analysis wherein, suppression of D genome was significantly less frequent in multiple tissue (Ramirez-Gonzalez et al., 2018). Earlier, homoeolog expression was also studied for wheat under infection with *Fusarium pseudograminearum*, wherein the expression bias was observed for B and D subgenomes (Powell, et al., 2017). Utilizing RNAseq data, changes in the pattern of the homoeologous gene expression were also reported in bread wheat (Leach et al., 2014). Therefore, such studies are important to pin-point actively expressed/induced homoeologs that could be targets for genetic improvement. However, the genome biasness and its association with the secretion of siderophores, genotypic variability in a complex wheat genome has yet to be investigated. Previous studies have shown that the phytosiderophores release mechanism in wheat is effective when the three genomes A, B and D contribute synergistically to induce its biosynthesis and release. In general, genome induction biasness for A and B genome was observed for the hexaploid-wheat under Fe stress (Fig. 3B). Interestingly, these genomes were also suggested to be important during starvation of other micronutrients like Zn (Tolay et. al., 2001). These observations provide clue for the importance of A and B genomes during micronutrient starvation in hexaploid wheat when compared to their progenitor. The PS release intensity is also dependent on complementary action of the genotypic ploidy with the following the pattern, *T. aestivum* (BBAADD) > *T. dicoccum* (BBAA) > *T. monococcum* (AA) > *Ae. tauschii* (DD) (Ma et al., 1999; Tolay et al., 2001).

Multiple genes for the salvage pathway were highly up-regulated including *NAS1, NAS2* and *NAAT1* (Supplementary Table S3 and Supplementary Table S5). The contribution of these transcripts arises from different genomes suggesting that all the genomes of wheat are capable of responding to the starvation condition. Based on our RNAseq analysis it is likely that the gene encoding for *TaYS1A* and *TaYSL9* could be the putative transporter for Fe-siderophore complex. Previous, studies have indicated the high response of *TaYS1A* under Fe starvation in roots and shoots (Kumar et al., 2019). Out of the closest orthologs of wheat-*YS1A, HvYS1A* and *ZmYS1A, HvYS1A* has been shown to be involved in specific transport of Fe whereas, ZmYS1 was reported to have broad substrate specificity (Schaaf et al., 2004). Our RNAseq data also revealed a conserved function of genes from monocots and dicots during the Fe regulation and known important Fe regulators such as PYE and BRUTUS also showed differential expression in wheat roots (Ivanov et al., 2012). Same for TFs spanning multiple gene families such as MYB (Rubio et al., 2001), *bHLH* (Colangelo et al., 2004; Jakoby et al., 2004; Ogo et al., 2007), *C2H2, NAC, AP2-ERB* (Kim et al., 2012), and *WRKY* (Devaiah et al., 2007) were characterized in Arabidopsis and rice for their involvement in nutrient uptake which were found to be highly represented in root of Fe-starved wheat. Similarly, hemerythrin motif-containing protein were also overexpressed in our study. As in case of other graminaceous plants like rice, hemerythrin domain containing proteins including OsHRZ homolog for Arabidopsis BRUTUS was also differentially perturbed in – Fe (Kobayashi et al., 2014). Overall, based on the information obtained from our studies future research focus should be able to gain better insights of the molecular functions of these genes using tilling population in wheat and/or a heterologous system.

Changes in metabolite content in response to Fe-deficiency can lead the plant to undergo adaptive processes to maintain iron homeostasis (Schmidt et al., 2014). A schematic representation of data provides a comprehensive summary of the metabolic and transcriptional changes that occur during Fe starvation in wheat roots (Fig. 9). The mobilization and transport of Fe inside the plant tissue depends on its speciation and complexation form. Depending on the tissue, Fe could form a complex with multiple metabolites such as nicotianamine, malate, citrate that facilitates the transport through the phloem and xylem (Grillet et al., 2014; Palmer et al., 2013). Our analysis for the metabolome of roots suggested predominance of additional organic compounds such as fumarate, acetate, glycerate etc. thereby, suggesting that they could be also involved in Fe-complexation to assist in long distance transport and remobilization to different tissues (Fig. 9). These metabolites are reported for their ability to complex with Fe(III) state during the transport in the xylem vessels as detected during the X-ray absorption near-edge structure spectroscopy based studies (Terzano, et al., 2013). Similarly, the accumulation of gylcerate could be responsible for added thrust towards glycerate-serine pathway during Fe starvation (Figure 9). Glycerate-serine pathway are important to generate methionine and precursor (S-adenosyl methionine) of DMA biosynthesis. Consistent with the metabolome data, the transcripts encoding these proteins were highly upregulated in our study as presented by the bins (Figure 9). Members of ABC-B transporters were also reported to participate in oxidative stress that was linked with metal stress including Fe starvation (Kispal et al., 1997; Schaedler et al., 2014). During our analysis we have identified a TAP type, ABCB subfamily transporter referred to as *TaABCB25* that was highly upregulated during Fe starvation (Fig. 4). The closest ortholog for wheat ABCB25 transporters were identified in rice (OsABCB25) and Arabidopsis (AtABCB27) (Supplementary Fig. S5A). The temporal-induction of *TaABCC25* was also re-confirmed in roots subjected to Fe starvation (Supplementary Fig. S5B). *Arabidopsis thaliana* ABCB27 was shown to have the ability to transport glutathione conjugate metabolite precursors for the Fe-S cluster assembly (Schaedler et al., 2014). Based on the presence of domain structure these ABCB transporters from rice, Arabidopsis and wheat qualify as half-size transporter (Supplementary Fig. S5C). In graminaceous species, glutathione related activity was observed to be highly upregulated under Fe deficient condition supporting our observation in the current study (Bashir et al., 2007). Enhanced GST activity in wheat under Fe starvation and the abundance of *TaABCB25* in roots, warrants further study of this gene for its role in mobilizing micronutrient uptake. Glutathione and GST plays an essential role during Fe starvation related responses in *Arabidopsis* and also quench reactive molecules to protect the cell from oxidative damage and prevent chlorophyll loss (Ramirez et al., 2013; Shanmugam et al., 2015; Kumar and Trivedi, 2018). Our analysis also suggests that high transcript and enzyme activity of GST could be linked with the primary metabolism since it accumulates fumarate (Fig. 9). In *Arabidopsis*, it has been shown that GST catalyses Glutathione-dependent isomerization of maleyl-acetoacetate into fumaryl-acetoacetate eventually leading to accumulation of fumarate and acetoacetate (Dixon et al., 2002). Additionally, GST is also linked with the conjugation of the reduced glutathione (GSH) forms to various substrate to make it water soluble. The sulphur derived metabolite and glutathione are well known to be involved in biosynthesis methionine, a well-known precursor for DMA biosynthesis (Forieri et al., 2013). Altogether, this supports the notion that in wheat role of glutathione is important during Fe starvation and ABC transporters might play a significant role in mobilization of metabolite/organic acid or specific siderophores.

In conclusion, in this study, the core components for Fe starvation response in hexaploid wheat have been identified so as to provide a better understanding of the molecular events that participate during Fe starvation in wheat roots. In particular, we provide line of evidence for the role of GST in the response to –Fe in roots. Complementary approaches, analytical and transcriptome analysis reinforce the importance of primary metabolism for reprogramming organic acids and amino acids thereby leading to the Fe homeostasis in wheat. The information here not only help to design strategies to improve plant response to Fe starvation in wheat, but will also foster Fe uptake and accumulation studies, which are required to boost productivity and grain nutritional quality through Fe biofortification programs.

## Acknowledgement

Authors thank Executive Director, NABI for facilities and support. This research was funded by the NABI-CORE grant to AKP and partial support from DST-SERB grant (PDF/2016/001355) to PG. GK and AK acknowledge NABI-SRF Fellowships. Technical help from Jagdeep Singh for the GC-MS analysis is highly appreciated. DBT-eLibrary Consortium (DeLCON) is acknowledged for providing timely support and access to e-resources for this work. The wheat genome resources developed by International Wheat Genome Sequencing Consortium are highly appreciated. We also acknowledge the critical comments given by Dr. Kerry Pedley, USDA, USA. We would like to thank Abhijeet Panwar (CDAC-Pune) for his help in the preparation of the Sankey plot.

## Supporting information

**Supplementary Fig. S1:** qRT-PCR validation of selected genes from the DEGs during Fe-deficient roots after 20 days of starvation. A total of 2 µg of RNA (DNA free) was used for cDNA synthesis and qRT-PCR was performed using gene specific primers (Supplementary Table S1). C_t_ values were normalized against wheat *ARF1* as an internal control.

**Supplementary Fig. S2:** Overview of genes modulated by the Fe starvation in wheat roots. MapMan overview demonstrating differentially expressed transcripts under iron starvation in general metabolic pathways. Log2 fold change values of DEGs were imported into MapMan. Read and blue bins represent up-regulation and down-regulation, respectively, in terms of log2 fold change, as shown by the scale.

**Supplementary Fig. S3:** MapMan visualization depicting the differentially expressed transcription factors families, with red and blue colored bins for up and down-regulated transcripts, respectively. Numbers in the scale represent fold changes in expression levels expressed as Log2.

**Supplementary Fig. S4:** Estimation of Nitrate levels under Fe-starvation using salicylic acid method. Three Biological replicate root samples of 10, 15 and 20 days after starvation (DAS) were completely dried for the extraction. Yellow coloration (in test tubes: lower panel) represents level of nitrate in the sample. Potassium Nitrate was used as standard in 0-70 µg concentrations.

**Supplementary Fig. S5:** Characterization of wheat ABCB25 transporter. (A) Phylogeny analysis of TaABCB25 along with its closest orthologs from rice and arabidopsis. (B) Expression analysis of *TaABCC25* in roots of wheat seedlings subjected to Fe starvation. Wheat seedlings (5-7 days old) were subjected to Fe stress and samples were harvested after 12 hours, 3, 6, 9 and 15 days(d) post starvation. The relative qRT-PCR was performed using wheat ADP-ribosylation factor (ARF) as an internal control gene. Fold accumulation was calculated with respect to the control roots. (C) Schematic comparision of different domains of AtABCB27, OsABCB25 and TaABC25. The number indicates the predicted amino acid position of the TM domains.

**Supplementary Table S1: List of primers used in the current study.** *wheat genes named according to rice RAP-DB/RefSeq based on KOBAS annotation.

**Supplementary Table S2: Summary of filtered and mapped reads for each sample.** Obtained RNA-seq reads were quality filtered using Trimmomatic v0.35. TopHat was used to map the obtained reads to the wheat genome (TGACv1).

**Supplementary Table S3: DEGs in response to Fe starvation in wheat roots** List of up-regulated genes, downregulated genes (sheet 2), genes exclusively expressed in response to Fe starvation (sheet 3). Table enlists Control and Fe starvation expression values, logFC for Fe starvation wrt Control samples. Each DEG is annotated with information like rice ortholog, gene definition, KEGG Orthology, Pathways and Pfam domains, which were obtained through KOBAS 3.0 stand-alone tool, using *Oryza sativa* RAP-DB and RefSeq as reference.

**Supplementary Table S4: List of 8473 homoeolog triads that were used for homoeolog induction and expression biasness analysis.** 8473 homoeolog triplets that were expressed in A, B as well as, D genome were selected for homoeolog specific analysis after filtering all the homoeolog triads obtained from from ensembl.

**Supplementary Table S5: Expression profiles of genes/gene families involved in Strategy-II mode of Fe uptake.** Strategy-II components of Fe uptake were identified and classified based on screening wheat genes annotated by KOBAS for respective KO IDs for NAS, NAAT, YSL and DMAS genes.

**Supplementary Table S6: Expression profiles of genes/gene families involved in Strategy-I mode of Fe uptake.** Strategy-I components of Fe uptake were identified and classified based on screening wheat genes annotated by KOBAS for respective KO IDs for AHA, IRT, FRO, PEZ genes.

**Supplementary Table S7: Gene Ontology analysis of up- and down-regulated genes in response to Fe starvation.** WEGO tool was used to categorize DEGs into GO categories and identify significant GO terms. Table lists the number and percentage of up- and down-regulated genes as well as p-values for each GO term.

**Supplementary Table S8: Expression profiles of genes/gene of different transcription factors those are differentially up- and down regulated.** MapMan was used to identify TFs and categorize them into TF families. Table gives logFC value for starvation *vs* control for each TF showing significantly altered expression. A gradient of red and green is used for up-regulated and down-regulated TFs respectively.

**Supplementary Table S9: Expression profiles of genes involved during the process of glutathione mediated detoxification process.** DEGs mapped to glutathione metabolism were identified using KOBAS. Table lists expression values and annotation for genes. Red color denotes up-regulated genes and green color represent down-regulation.

**Supplementary Table S10: GC-MS analysis of wheat roots subjected to 20 days of Fe starvation.** Each metabolite is represented with concentrations in three independent replicate manner. For concentration calculation, Individual metabolite area was normalized to sample weight and area of internal control (sorbitol). Metabolites with no detectable area in any of the conditions were considered to be the metabolite with minimum area. Delta method approximation was used to calculate standard errors (se) of log-ratio, se log-ratio = 1/ln 2√[(SET /T)2 + (SEC /C)2], where SET and SEC are standard errors of average –Fe and +Fe metabolite abundances. Metabolites with significant (p-value <0.05) differential abundance were plotted.

