## Supplemental Figures for "Integrative analysis of hexaploid wheat roots identifies signature components during iron starvation"

### Slide 1
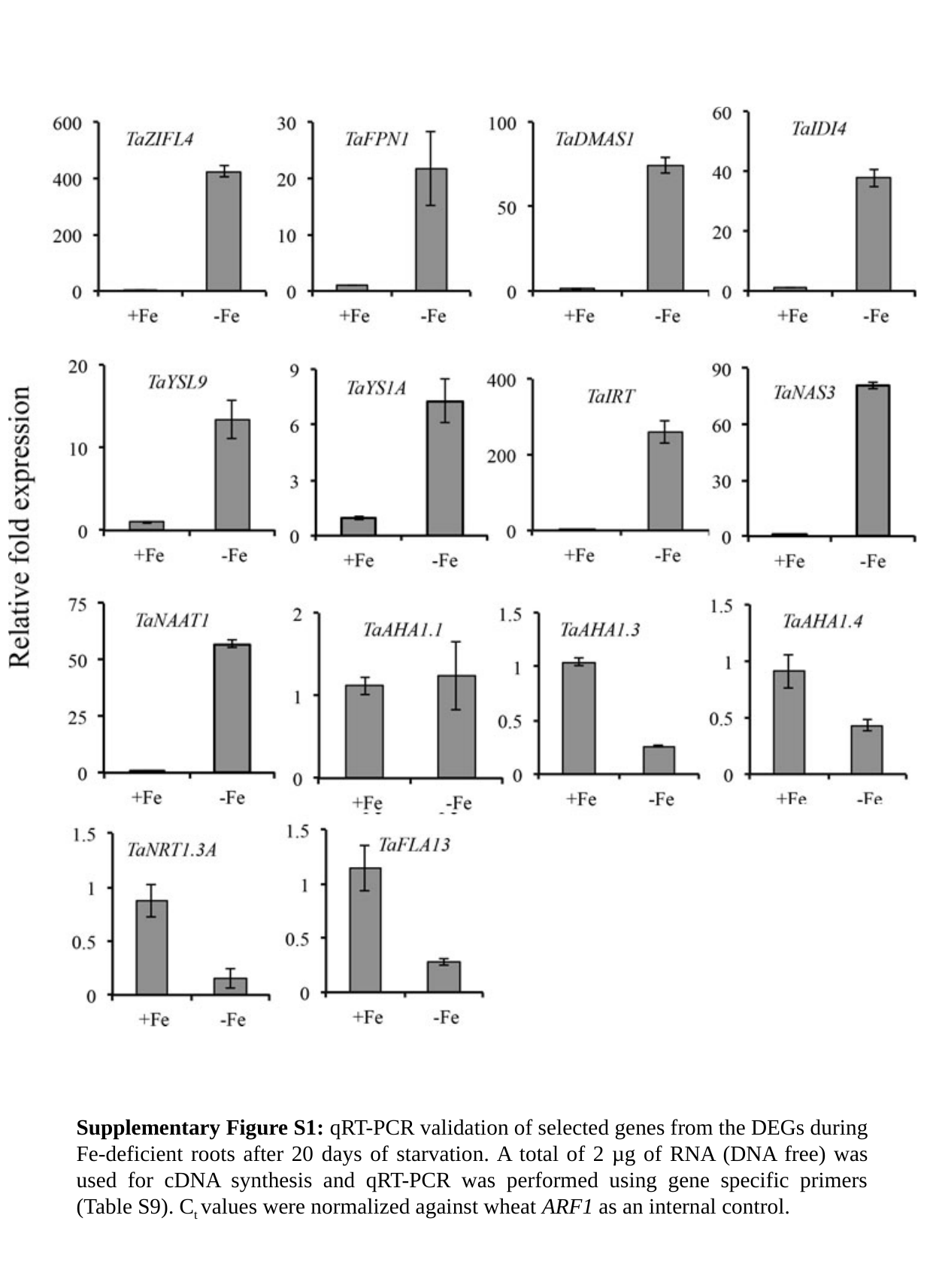

Supplementary Figure S1: qRT-PCR validation of selected genes from the DEGs during Fe-deficient roots after 20 days of starvation. A total of 2 µg of RNA (DNA free) was used for cDNA synthesis and qRT-PCR was performed using gene specific primers (Table S9). C_t_ values were normalized against wheat ARF1 as an internal control.

### Slide 2
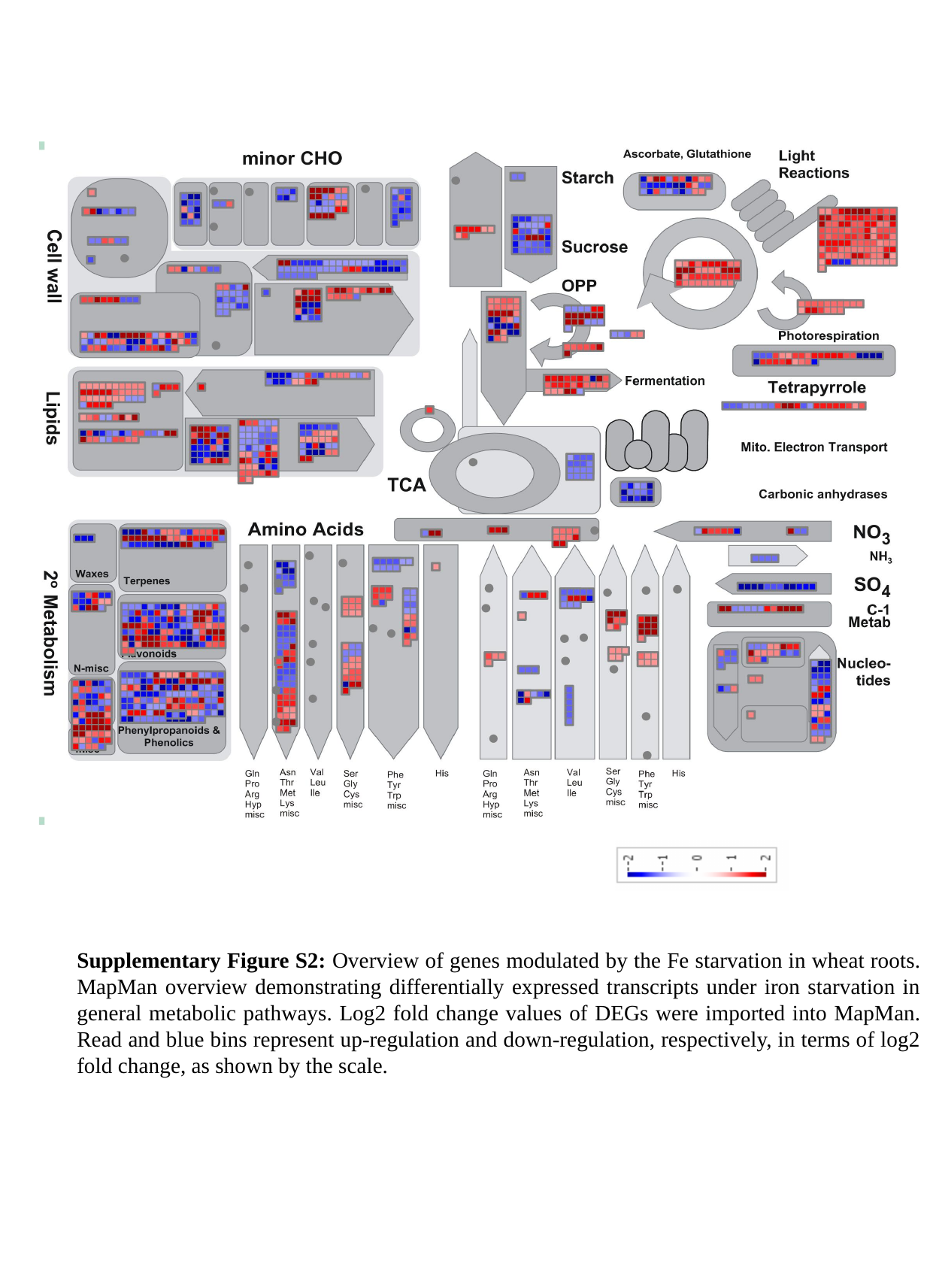

Supplementary Figure S2: Overview of genes modulated by the Fe starvation in wheat roots. MapMan overview demonstrating differentially expressed transcripts under iron starvation in general metabolic pathways. Log2 fold change values of DEGs were imported into MapMan. Read and blue bins represent up-regulation and down-regulation, respectively, in terms of log2 fold change, as shown by the scale.

### Slide 3
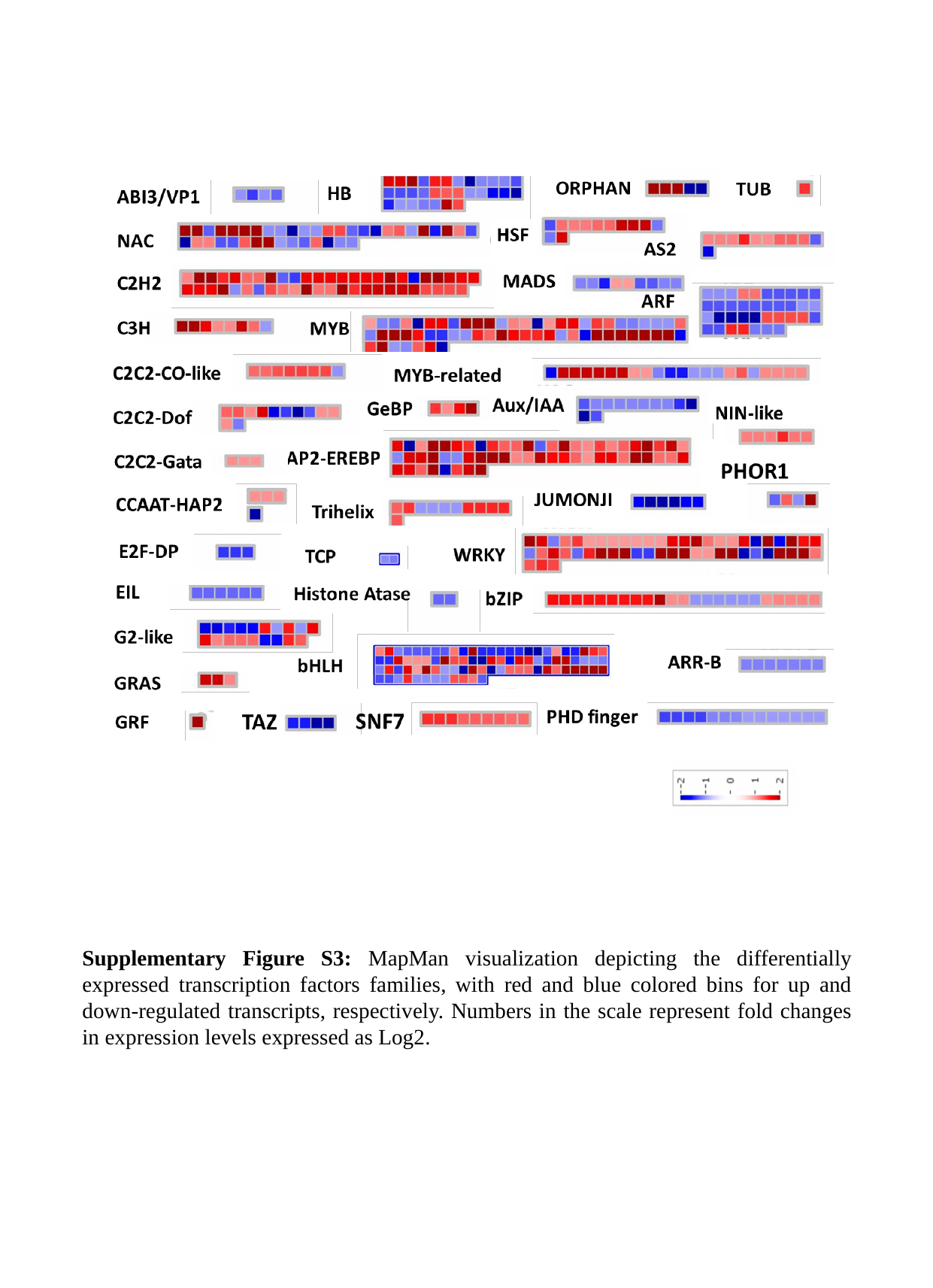

Supplementary Figure S3: MapMan visualization depicting the differentially expressed transcription factors families, with red and blue colored bins for up and down-regulated transcripts, respectively. Numbers in the scale represent fold changes in expression levels expressed as Log2.

### Slide 4
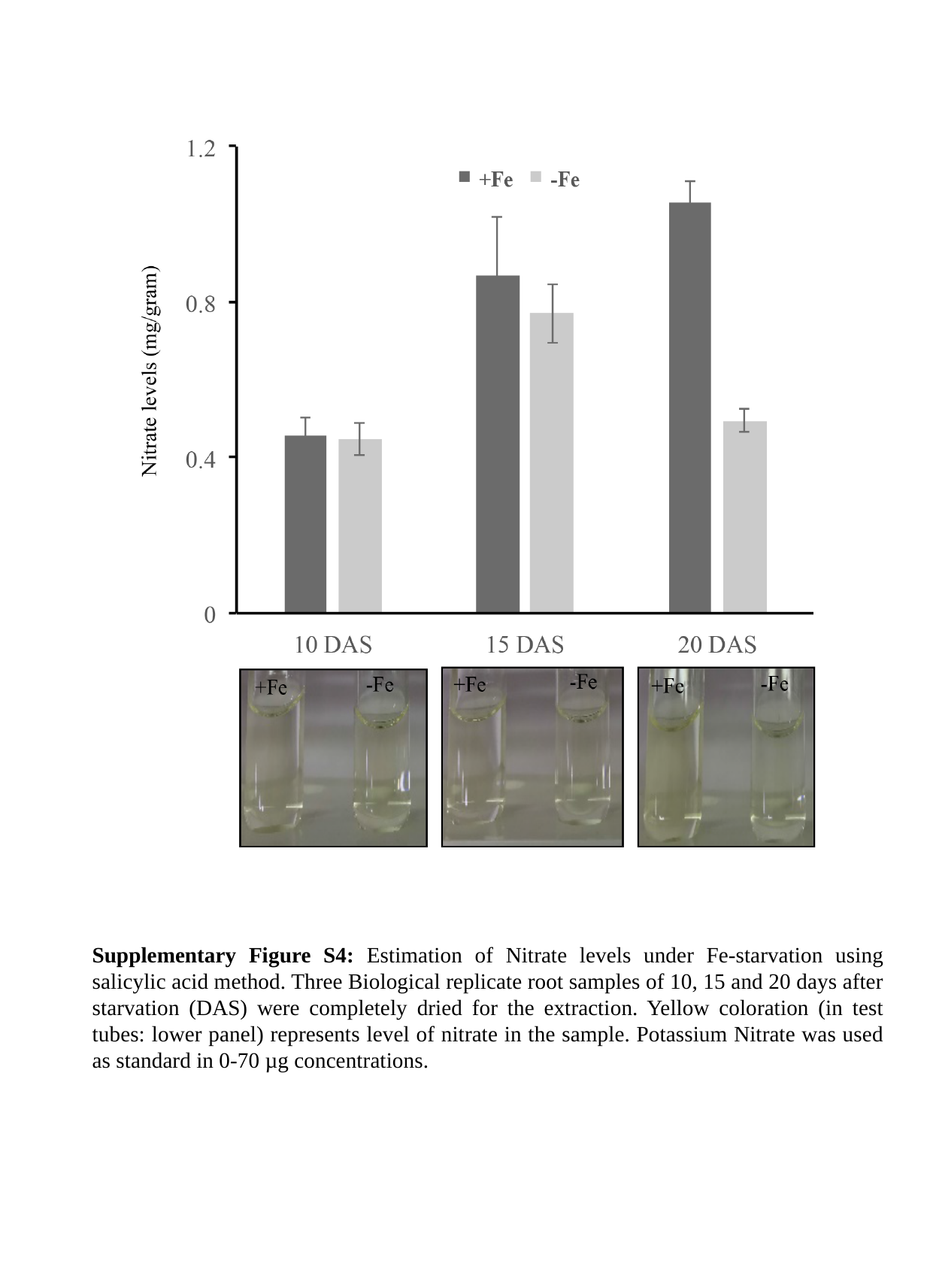

Supplementary Figure S4: Estimation of Nitrate levels under Fe-starvation using salicylic acid method. Three Biological replicate root samples of 10, 15 and 20 days after starvation (DAS) were completely dried for the extraction. Yellow coloration (in test tubes: lower panel) represents level of nitrate in the sample. Potassium Nitrate was used as standard in 0-70 µg concentrations.

### Slide 5
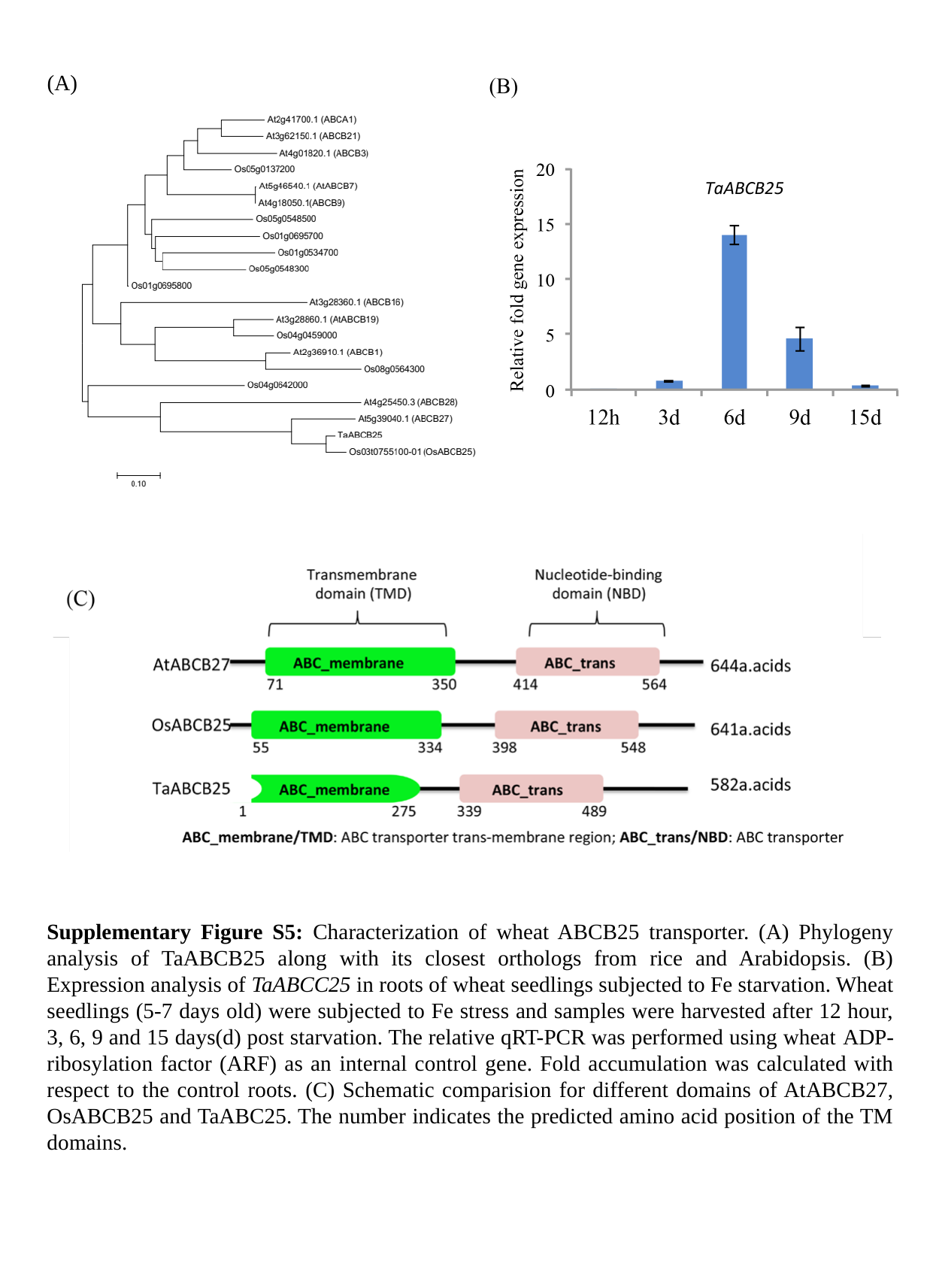

(A)
Supplementary Figure S5: Characterization of wheat ABCB25 transporter. (A) Phylogeny analysis of TaABCB25 along with its closest orthologs from rice and Arabidopsis. (B) Expression analysis of TaABCC25 in roots of wheat seedlings subjected to Fe starvation. Wheat seedlings (5-7 days old) were subjected to Fe stress and samples were harvested after 12 hour, 3, 6, 9 and 15 days(d) post starvation. The relative qRT-PCR was performed using wheat ADP-ribosylation factor (ARF) as an internal control gene. Fold accumulation was calculated with respect to the control roots. (C) Schematic comparision for different domains of AtABCB27, OsABCB25 and TaABC25. The number indicates the predicted amino acid position of the TM domains.
