## Supplementary material for "Integrative analysis of hexaploid wheat roots identifies signature components during iron starvation": Table S1

**Table S1: List of primers used in the current study.** **wheat genes named according to rice RAP-DB/RefSeq based on KOBAS annotation.*

| Gene name | Primer sequence 5’-3’ | Amplicon size in bp |
| --- | --- | --- |
| *TaARF1* | F: TGATAGGGAACGTGTTGTTGAGGC | 234 |
|  | R:AGCCAGTCAAGACCCTCGTACAAC |  |
| *TaDMAS1* | F:CGACATGCGGGCCGTGTGGGAG | 148 |
|  | R:CACCTCCACCTGATTGACGGTGGG |  |
| *TaPTR2-B^*^* | F:CGACGCCTGGCTCGGCCGCT | 151 |
|  | R:CGCCACCAGGTAGAGCGCGACG |  |
| *TaYSL9* | F:CGGGTCCTACCTGCTCGGGCTC | 156 |
|  | R:CTGAGAGGGACAAGCGCGAGAATCC |  |
| *TaYS1A* | F:ATGACATACGTTGGTGCCGGGATGATTTGCCC | 202 |
|  | R:CCCCCATGATGAGAGCTATGCATATGAAGGCC | |
| *TaFPN1^*^* | F:GCGCTCGCGGCGCTCTCAACTC | 142 |
|  | R:GCAGCTTGCAGCTCAGGTCGATCC |  |
| *TaIDI4^*^* | F:CGACAACGGCGCCGCCAAGCC | 154 |
|  | R:GCCATCAAAATTGGGAAAGCCCTGACC |  |
| *TaNRT2.2^*^* | F:GTCATCGCCGAGTACTACTTCGAC | 147 |
|  | R:ACGCATGCCGAAGTAGCGGGC |  |
| *TaNRT2.3^*^* | F:GCGCTTCTTCACGGGCTTCTCGC | 150 |
|  | R:GGCATGATGAACTGCACGGCGCC |  |
| *TaCytP450^*^* | F:GGCTCTTCGAGAACTGGTGGCCG | 141 |
|  | R:GGACCAGCCTGAGGTTCTCGGG |  |
| *TaNRT1.3A^*^* | F:GGCCTCTGCTCTGGCGCCATG | 156 |
|  | R:GGTGAGCGGCTGCTTGAGGCC |  |
| *TaFLA 13^*^* | F:CGGTCGCTCATGCTGCACCACG | 152 |
|  | R:GTGGCCCAGCCCGACACGACG |  |
| *TaCAD3^*^* | F:CCTACAACTCGGTCGACCGTGACG | 152 |
|  | R:GCTGTACACGGTGATGCCGGCG |  |
| *TaFRO^*^* | F:GTGGCACGCGGCTCGTCGCTC | 121 |
|  | R:CGTAGCAGAGCCCATGAGCGGTG |  |
| *TaIRT1.1^*^* | F:CGACTCGCTCATGCTCACCTTCTAC | 191 |
|  | R:GCTGCATCTGGCCGGCCTCAG |  |
| *TaAHA1.1^*^* | F:GCAATCGAGGAGATGGCGGGCATGG | 217 |
|  | R:CCTCTTTCGGATCAGCTAGCATCCC |  |
| *TaAHA1.2^*^* | F:GCGAGCAGGAGGCATCAATCTTGG | 162 |
|  | R:CATCCCCAGGGTTCTTGGTCACC |  |
| *TaAHA1.3^*^* | F:GGAGGCATTCCCATAGCGATGCCCA | 134 |
|  | R:CAAAGAACATCCATGCCGGCCATCTC |  |
| *TaAHA1.4^*^* | F:CGGCGGCATCCCGATCGCGAT | 136 |
|  | R:GCAGAGCACGTCCATGCCGGCC |  |
| *TaZIFL4^*^* | F:GATTCAAGAGCTGCTCCCCATAGAGAAGTT | 217 |
|  | R:CACCTGAAACTGCAAGAACTTGGCCA |  |
| *TaNAS3* | F:TACGAGCTCCTGGCGCGCTA | 176 |
|  | R:AACAGCTTGCTGGCGCGGTC |  |
| *TaNAAT1* | F:ATCGGCCGACGACATCTTCCTCA | 156 |
|  | R:CGAACCTCCAGCTTGTTGAACGC |  |
| *TaABCB25^*^* | F:ACAAGACTGTCGTGGGAGAG | 168 |
|  | R: ACGAGTCCATTGCATCCTGA |  |

- Orthology based naming of *Triticum aestivum* genes according to KOBAS annotation based on *Oryza sativa* RAP-DB/RefSeq.
